# Conformational dynamics regulate SHANK3 actin and Rap1 binding

**DOI:** 10.1101/2020.11.12.379222

**Authors:** Siiri I Salomaa, Mitro Miihkinen, Elena Kremneva, Ilkka Paatero, Johanna Lilja, Guillaume Jacquemet, Joni Vuorio, Lina Antenucci, Fatemeh Hassani-Nia, Patrik Hollos, Aleksi Isomursu, Ilpo Vattulainen, Eleanor T. Coffey, Hans-Jürgen Kreienkamp, Pekka Lappalainen, Johanna Ivaska

## Abstract

Actin-rich cellular protrusions direct versatile biological processes from cancer cell invasion to dendritic spine development. The stability, morphology and specific biological function of these protrusions are regulated by crosstalk between three main signaling axes: integrins, actin regulators and small GTPases. SHANK3 is a multifunctional scaffold protein, interacting with several actin-binding proteins, and a well-established autism risk gene. Recently, SHANK3 was demonstrated to sequester integrin-activating small GTPases Rap1 and R-Ras to inhibit integrin activity via its N-terminal SPN domain. Here, we demonstrate that SHANK3 interacts directly with actin using its SPN domain. Actin binding can be inhibited by an intramolecular closed conformation of SHANK3, where the adjacent ARR domain covers the actin-binding interface of the SPN domain. Actin and Rap1 compete with each other for binding to SHANK3 and loss of SHANK3-actin binding augments inhibition of Rap1-mediated integrin activity. This dynamic crosstalk has functional implications for filopodia formation in cancer cells, dendritic spine morphology in neurons and autism-linked phenotypes *in vivo*.

## Introduction

The distinct cell types of a living organism can adopt remarkably versatile shapes that are dynamically regulated during physiological processes. Short-lived actin-rich cell protrusions such as filopodia, membrane ruffles and lamellipodia as well as more stable structures such as dendritic spines, which mature from filopodia-like structures, are important contributors to cell shape and functionality ^1,2^. In adherent cells, these structures receive input from several sources including regulators of the actin cytoskeleton, integrin-mediated cell-extracellular matrix interactions and small GTPase signaling ^3–5^. Thus, crosstalk between these signals must be somehow carefully balanced within a cell.

SHANK3 is a structural scaffold protein predominantly studied in the post-synaptic density (PSD) of neurons owing to its well-established role in autism-spectrum disorders (ASD). SHANK3 mutations are associated with ASD ^6–9^, and SHANK3 epigenetic dysregulation has been reported in up to ~15% of ASD cases ^10^. In addition, SHANK3 mutations are linked to schizophrenia and Phelan-McDermid syndrome (22q13 deletion syndrome) highlighting the importance of SHANK3 in neuronal development ^6,8,11–13^. SHANK3 is also widely expressed outside of the central nervous system ^14,15^ with largely unknown functions. Unbiased RNAi screening in multiple cancer cell types and normal cells ^14,16^ revealed that SHANK3 inhibits integrin-mediated cell adhesion. The N-terminal SPN domain of the protein adopts an unexpected Ras-association (RA) domain-like fold that binds and sequesters active Rap1 GTPase with high affinity, preventing recruitment of the integrin activator protein talin, and attenuating integrin function ^14^. Importantly, Rap1 binding ^14^, and the associated inhibition of integrin activity were the first indication of a biological function for the evolutionally conserved SHANK3 SPN domain ^17^. Interestingly, two autism-linked SHANK3 patient mutations, R12C and L68P ^8^, are within the SHANK3 SPN domain, and impair the ability of SHANK3 to bind to Rap1 and inhibit integrin activation ^14^.

In neurons, SHANK3 associates with different actin regulators including Abi1 ^18,19^, Abp1 ^20^, α-fodrin 21, SHARPIN ^22^, βPIX ^23^, CaMKKIIα ^24^, IRSp53 ^25^, and cortactin ^26,27^. In the context of ASD, SHANK3 mutations contribute to disease pathogenesis through dysregulation of the actin cytoskeleton ^2,8,28–30^ and ASD symptoms of *Shank3-*deficient mice are alleviated by targeting actin regulators ^30^. Thus, SHANK3 regulation of actin dynamics is required for normal neuronal development and function. However, there are no reports of SHANK3 interacting directly with actin and the potential role of SHANK3 in regulation of the actin cytoskeleton has not been investigated in non-neuronal cells.

Here, we present evidence that SHANK3 binds actin directly through its N-terminal SPN domain, and that this interaction is subject to regulation through the formation of an auto-inhibited SHANK3 conformation. Furthermore, we show through molecular simulations that active Rap1 may stabilize the closed conformation, and actin and active Rap1 compete for binding to the SPN domain. Therefore, the balance between actin and Rap1 binding to SHANK3 controls integrin activity and integrin-dependent filopodia in cancer cells, dendritic spine morphology in neurons, and impacts SHANK3 functionality in an ASD zebrafish model.

## Results

### The SHANK3 SPN domain inhibits filopodia formation and colocalizes with actin

To gain insight into the role of SHANK3 in regulating actin-rich plasma membrane protrusions in non-neuronal cells, we imaged filopodia, cellular protrusions that are dependent on integrin activity and which are implicated in cancer cell invasion ^31–33^. Filopodia were induced by expressing the fluorescently tagged motor-protein myosin-X (MYO10) in U2OS (human osteosarcoma) cells ^33,34^ and the dependence on integrin activity was validated by co-expressing known integrin activators, talin-1 and kindlin-2, which significantly increased the number of MYO10-positive filopodia (Extended Data Fig. 1A-B). Expression of full-length GFP SHANK3 (functional domains highlighted in Fig. 1A) reduced the number of MYO10-positive filopodia significantly (Fig. 1B-C) and the effect was more prominent with the GFP SHANK3 SPN domain alone (from here on referred to as GFP SPN, Fig. 1D-E), which interacts with Rap1-GTP, and is sufficient to inhibit integrins ^14^.

**Figure 1.**
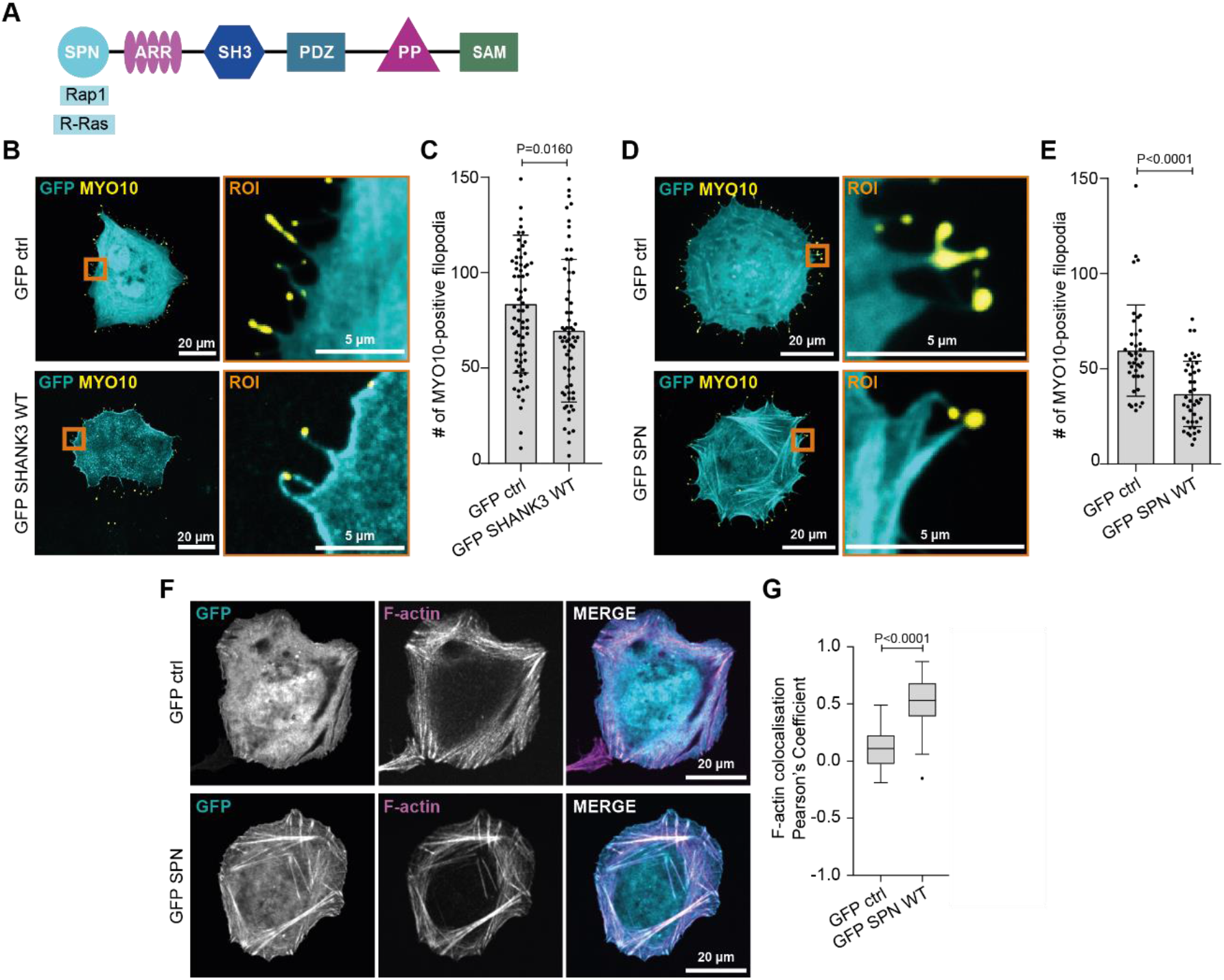
The SHANK3 SPN domain inhibits MYO10-positive filopodia formation and colocalizes with F-actin. **A,** Schematic of SHANK3 functional domains. SPN, Shank/ProSAP N-terminal domain; ARR, ankyrin repeat region; SH3, Src homology 3 domain; PDZ, PSD-95/Discs large/ZO-1 domain; PP, proline-rich region; SAM, sterile alpha motif domain. Known direct binding partners of the SPN domain are indicated. **B, C,** Analysis of filopodia formation in U2OS cells co-expressing either GFP control or GFP SHANK3 together with MYO10-mCherry (to induce and visualize filopodia) and plated on fibronectin for 2 h. Representative bottom plane confocal images (B) and quantification of filopodia numbers (C) are shown. **D, E,** Analysis of filopodia formation in U2OS cells co-expressing either GFP control or GFP SPN together with MYO10-mCherry and plated on fibronectin for 2 h. Representative bottom plane confocal images (D) and quantification of filopodia numbers (E) are shown. **F, G,** Analysis of F-actin (phalloidin-647) and GFP localization in U2OS cells expressing either GFP control or GFP SPN and plated on fibronectin (3-4 h). Representative bottom plane confocal images (F) and quantification using the coloc2 ImageJ plugin (G) are shown. Orange squares highlight regions of interest (ROI), which are magnified. All representative images and data are from n = three independent experiments. Data are mean ± standard deviation (s.d.) (C, E) or presented as Tukey box plots with median and the interquartile range (IQR) (whiskers extend to 1.5x the IQR and outliers are displayed as individual points). Statistical analyses: (C, E) Mann-Whitney two-tailed T-test. (G) Kruskal-Wallis non-parametric test and Dunn’s multiple comparisons post hoc test. Number of cells analyzed: (C) 74 (GFP ctrl) and 67 (GFP SHANK3 WT). (E) 41 (GFP ctrl) and 43 (GFP SPN). (G) 79 (GFP ctrl) and 84 (GFP SPN).

Talin and kindlin colocalize with active integrins in filopodia tip adhesions ^33^ and talin supports integrin activation at the tip ^32,35^. In contrast, SHANK3 did not localize to filopodia tips (Extended Data Fig. 1C), suggesting an alternative mechanism of filopodia regulation such as limiting the availability of Rap1-GTP in the cell or regulating the actin cytoskeleton. Imaging revealed GFP SPN overlap with filament-like structures proximal to the base of filopodia (Fig. 1D), and colocalization with filamentous actin (F-actin) within the cell body in U2OS (Fig. 1F-G) and HEK293 (human embryonic kidney) cells (Extended Data Fig. 1D, E). These data imply that the SHANK3 SPN domain is recruited to actin filaments in cells.

### The SHANK3 SPN domain binds F-actin directly

SHANK3 is a large scaffold protein with six functional domains, which mediate interaction with many actin-associated and actin-binding proteins, including β-PIX ^23^, cortactin ^26,27^, Abi-1 ^19^, and Abp1 ^18^ (Extended Data Fig. 1F). However, there are no previous reports of a direct interaction between SHANK3 and F-actin and, as such, it is unclear whether SHANK3 could regulate actin directly in addition to facilitating the recruitment of actin regulators to actin. To investigate this, we studied the localization of the SHANK3 SPN domain and other SHANK3 fragments in U2OS cells stained for F-actin. While the SPN domain (residues 1-92) colocalized with F-actin (Fig. 1F-G, Extended Data Fig. 1D-E), similar overlap with filamentous structures was not observed in cells expressing longer SHANK3 fragments or full-length SHANK3 (Extended Data Fig. 1G). SHANK3 shuttles in and out of the nucleus, and deletion of the C-terminal region increases its nuclear accumulation ^36,37^. Furthermore, SHANK3 has three nuclear localization sequences (NLS) C-terminal to the ARR domain at positions 350-359, 459-468 and 655-686 ^38^. Accordingly, the longer SHANK3 constructs displayed a predominantly nuclear localization while the full-length SHANK3 localized throughout the cell (Extended Data Fig. 1G), indicating that the SPN domain localizes to actin in a manner that is somehow inhibited in the context of the full-length protein.

To test whether the SPN domain binds to non-muscle β/γ-actin directly, we employed an F-actin co-sedimentation assay with recombinant SPN protein and purified actin, and observed that GST SPN interacts with actin filaments (Extended Data Fig. 2A-B). To address whether SPN binding affects the dynamics of the actin filaments, we performed fluorometric actin filament disassembly assays in the presence and absence of cofilin-1. Addition of GST SPN did not alter spontaneous or cofilin-1-induced actin filament disassembly (Extended Data Fig. 2C-D). These data demonstrate that the SHANK3 SPN domain interacts directly with actin filaments without altering their stability.

### Identification of the SHANK3 SPN actin-binding site

The SHANK3 SPN domain is structurally similar to the N-terminal F0 domain of talin (Extended Data Fig. 2E) ^14,39^. The kindlin-2 F0 domain, which also adopts a similar fold (Extended Data Fig. 2F), binds actin directly ^40^. This prompted us to compare the structures of kindlin F0 and SHANK3 SPN in more detail. Superimposition of the SHANK3 SPN domain with the F0 domains of kindlin-1 and −2 revealed a corresponding spatial alignment between SPN residues Q37 and R38 and the kindlin F0 actin-binding residues L47 and K48 (Fig. 2A-B). Furthermore, the local charge distribution of the predicted binding sites correlated well between kindlin F0 and the SHANK3 SPN domain (Extended Data Fig. 2G). Thus, we hypothesized that Q37 and R38 residues in the SPN domain may contribute to actin binding (Fig. 2B). Replacing these residues with alanines (GFP SPN Q37A/R38A) abolished SPN co-localization with actin stress fibers in unconstrained cells and in cells plated on crossbow-shaped micropatterns (Fig. 2C-E). Interestingly, the GFP SPN R12C mutant, with compromised Rap1-binding ^14^, colocalized with actin stress fibers similarly to WT SPN (Fig. 2C-E), indicating that the interaction between the SPN domain and Rap1 is not required for SPN recruitment to actin filaments in cells.

**Figure 2.**
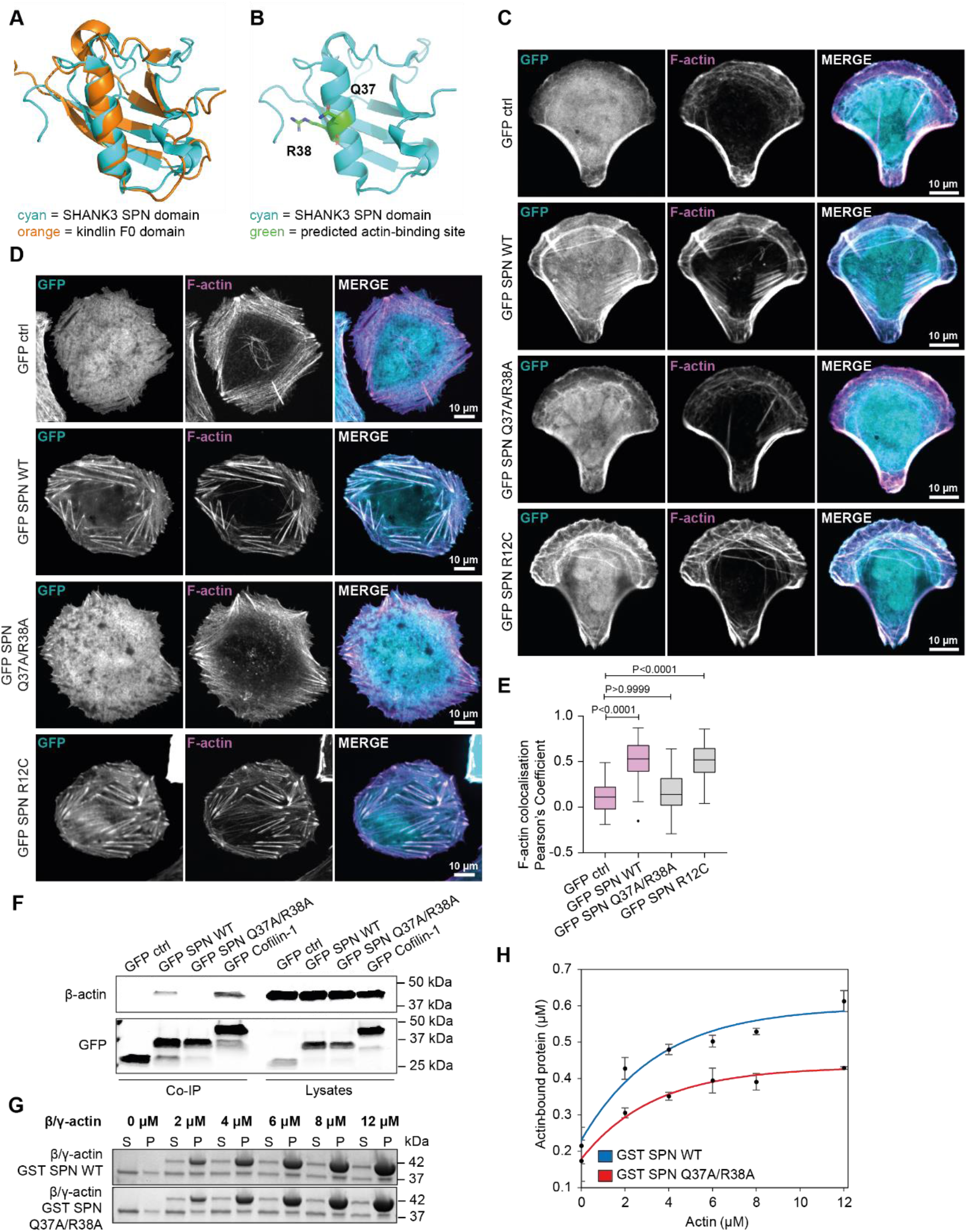
The SHANK3 SPN actin interaction is inhibited by mutation of the predicted actin-binding site. **A,** Superimposition of the SHANK3 SPN domain and the kindlin-1 F0 domain using Pymol (PDB codes: 5G4X, 2KMC). **B,** Structure of the SHANK3 SPN domain (5G4X) where actin-binding residues Q37/R38 are highlighted as green sticks. **C,** Analysis of F-actin (phalloidin-647) and GFP localization in U2OS cells expressing either GFP control or GFP-tagged SPN WT, SPN Q37A/R38A or SPN R12C and plated on fibronectin–coated crossbow-shaped micropatterns. Representative bottom plane confocal images are shown. **D, E,** Analysis of F-actin (phalloidin-647) and GFP colocalization in U2OS cells expressing either GFP control or GFP-tagged SPN WT, Q37A/R38A or R12C and plated on fibronectin-coated glass-bottom dishes (3-4 h). Representative bottom plane confocal images (D) and quantification (E) using the coloc2 ImageJ plugin are shown. Pink-colored boxes have been shown earlier in Figure 1G. **F,** GFP-trap pulldowns in U2OS cells expressing GFP control (negative control), GFP-tagged Cofilin-1 (positive control), GFP SPN WT or Q37A/R38A. Input lysates and immunoprecipitated (IP) samples were analyzed using β-actin and GFP antibodies as indicated. **G, H,** Analysis of GST-tagged SPN WT or SPN Q37A/R38A interaction with β/γ-actin filaments in co-sedimentation assays using SDS-PAGE, followed by Coomassie Blue staining. A representative co-sedimentation assay experiment (G) and quantification (H) where the amounts of co-sedimenting SPN WT and Q37A/R38A were plotted against actin concentration are shown. Please note that at high concentrations the amount of co-sedimenting GST SPN plateaued at ~0.65 μM, indicating that 30-40 % of GST SPN was inactive and unable to bind actin filaments. The Exponential equation *f* = *y*_*o*_ + *a* * (1 − *exp*^−*b*\**x*^) was applied to fit experimental data; n = 3 independent experiments; error bars represent s.e.m. S, supernatant fraction; P, pellet fraction. All representative micrographs, immunoblots and data are from n = three independent experiments. Data are presented as Tukey box plots (E) or as exponential curve with s.e.m. (H). Statistical analyses: (E) Kruskal-Wallis non-parametric test and Dunn’s multiple comparisons post hoc test. Number of cells analyzed: (E) 57 (GFP ctrl), 88 (GFP SPN WT), 90 (Q37A/R38A) and 68 (R12C).

GFP SPN Q37A/R38A was also defective in pulling down β-actin from cell lysates when compared to GFP SPN WT or GFP Cofilin-1 (positive control) (Fig. 2F). In addition, in actin co-sedimentation assays β/γ-actin filament binding to GST SPN Q37A/R38A was reduced compared to WT GST SPN (Fig. 2G-H, Extended Data Fig. 2A, H). Taken together, our findings indicate that the SHANK3 SPN domain interacts with F-actin through a similar mechanism to kindlin-2 F0 domain, and consequently the Q37A/R38A point mutant reduces SHANK3 SPN domain binding to actin filaments *in vitro* and in cells.

### Crosstalk between SHANK3 SPN-actin binding and integrin inhibition

The active-Rap1-binding interface of the SHANK3 SPN domain (including the conserved, ASD associated SPN R12 residue ^14^) is distinct from the SPN actin-binding site (Q37/R38) (Extended Data Fig. 3A). This suggests that the SHANK3 integrin inhibitory and actin-binding functions could be independent as they are mediated through different interfaces of the SPN domain. To test this hypothesis, we employed an established flow cytometry-based integrin activity assay where active integrin levels are determined as a ratio of ligand-bound integrins (labelled fibronectin fragment domains 7-10) over total cell-surface β1-integrins ^14^. We have earlier shown that expression of GFP SPN WT, but not GFP SPN R12C (Rap1-binding defective mutant), reduces integrin activation ^14^. Here, we observed that the actin-binding-deficient SPN Q37A/R38A mutant inhibited integrins significantly and more potently than GFP SPN WT (Fig. 3A). Thus, reduced actin binding augments the integrin-inhibiting function of the SHANK3 SPN domain, possibly due to increased availability of the SPN domain to bind to plasma membrane-localized Rap1-GTP.

**Figure 3.**
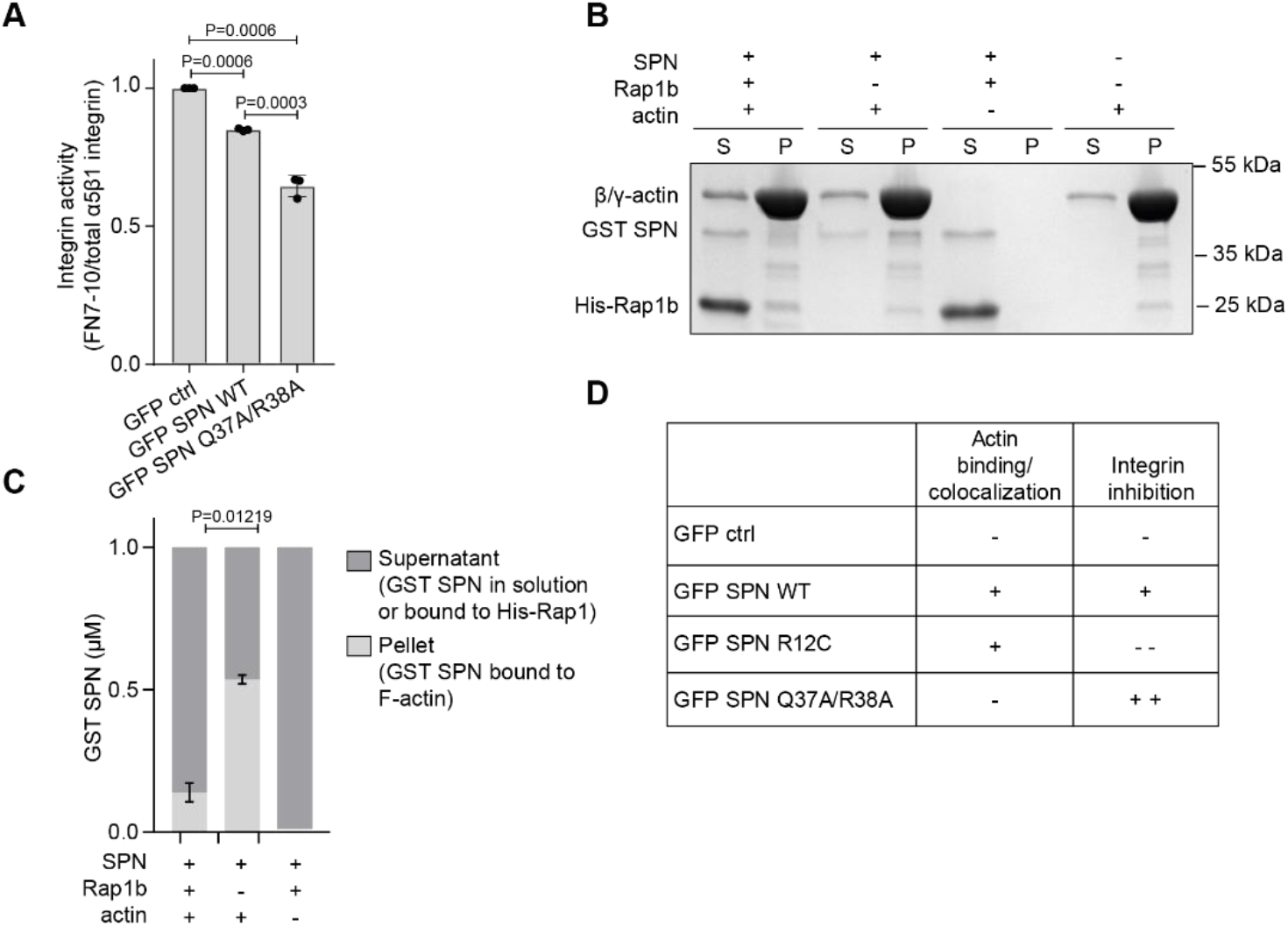
Loss of actin binding enhances SHANK3 SPN-mediated integrin inhibition. **A,** Flow cytometry analysis of integrin activity (fibronectin (FN) fragment FN 7-10 binding relative to total cell-surface α5β1-integrin) Quantification shows reduced active cell-surface integrin (FN 7-10 binding) in CHO cells expressing GFP-tagged SPN WT or SPN Q37A/R38A compared to GFP control. **B, C,** Analysis of GST SPN (1 μM) interaction with β/γ-actin filaments (12 μM) in the presence or absence of active GMPPCP-loaded (GTP-analogue) His-Rap1b (4 μM). A representative cosedimentation experiment analyzed by SDS-PAGE and Coomassie Blue staining (B) and quantification of the proportion of SPN in the pellet fraction (P, represents SPN bound to actin) versus the supernatant fraction (S, represents soluble protein not bound to actin) (C) are shown. The addition of active His-Rap1b increases the amount of GST-SPN remaining in the supernatant and not co-sedimenting with actin. Five independent experiments. **D,** Table summarizing the effects of different SPN mutants in actin binding and integrin inhibition. All immunoblots and data are from three independent experiments unless otherwise indicated. Data represent mean ± s.d., (A) or mean ± s.e.m (C). Statistical analysis: (A) Welch’s t-test with subsequent Bonferroni correction. (C) Mann-Whitney nonparametric test.

To test if active Rap1 binding to the SPN domain would influence SPN–actin interaction, we employed a modified version of the actin co-sedimentation assay. We incubated recombinant SPN domain with F-actin in the presence or absence of the active His-Rap1b loaded with a GTP-analogue (GMPPCP) (the SPN domain binds specifically to active Rap1 ^14,41^). In the absence of active Rap1, a large pool of the SPN domain co-sedimented with actin filaments (Fig. 3B-C). The addition of active Rap1 markedly decreased the amount SPN domain co-sedimenting with actin filaments (Fig. 3B-C) and a large proportion of the SPN domain remained in solution together with active Rap1 (Extended Data Fig. 3B). These data suggest that active Rap1-GTP and actin filaments compete with each other for binding to the SPN domain even though their binding sites on the SPN domain are distinct (Extended Data Fig. 3A, Fig. 3D) and can be independently disrupted by specific mutations.

### SPN-ARR fold opening dynamically regulates SPN‒actin interaction

Many adhesion and actin-regulating proteins, such as talin, formins, ezrin-radixin-and moesin (ERM) family proteins and N-WASP are autoinhibited by protein folding ^42–45^. As there was no clear overlap between the SPN-ARR fragment or full-length SHANK3 with actin filaments in cells (Extended Data Fig. 1G), we hypothesized that the conformation of SHANK3 may regulate its actin binding function. In the published crystal structure, the SPN-ARR fragment of SHANK3 ^14^ adopts a closed conformation that is mediated by intramolecular bonds between the SPN and ARR domains. In addition, in full-length SHANK3 the closed conformation inhibits binding of α-fodrin, SHARPIN and exogenous SPN to the ARR domain ^46^. The SPN actin-binding residues Q37 and R38 are located proximal to the SPN-ARR domain interface (Fig. 4A), and may therefore be inaccessible for actin binding when the fold is in a closed state. To test this hypothesis, we first analyzed recombinant SPN-ARR binding to actin. In contrast to the SPN domain alone, recombinant SPN-ARR did not co-sediment with filamentous β/γ-actin (Extended Data Fig. 4A-B) even though both SPN and SPN-ARR demonstrated effective binding to GFP Rap1 Q63E (active Rap1 mutant), confirming that both proteins were functional (Extended Data Fig. 4C).

**Figure 4.**
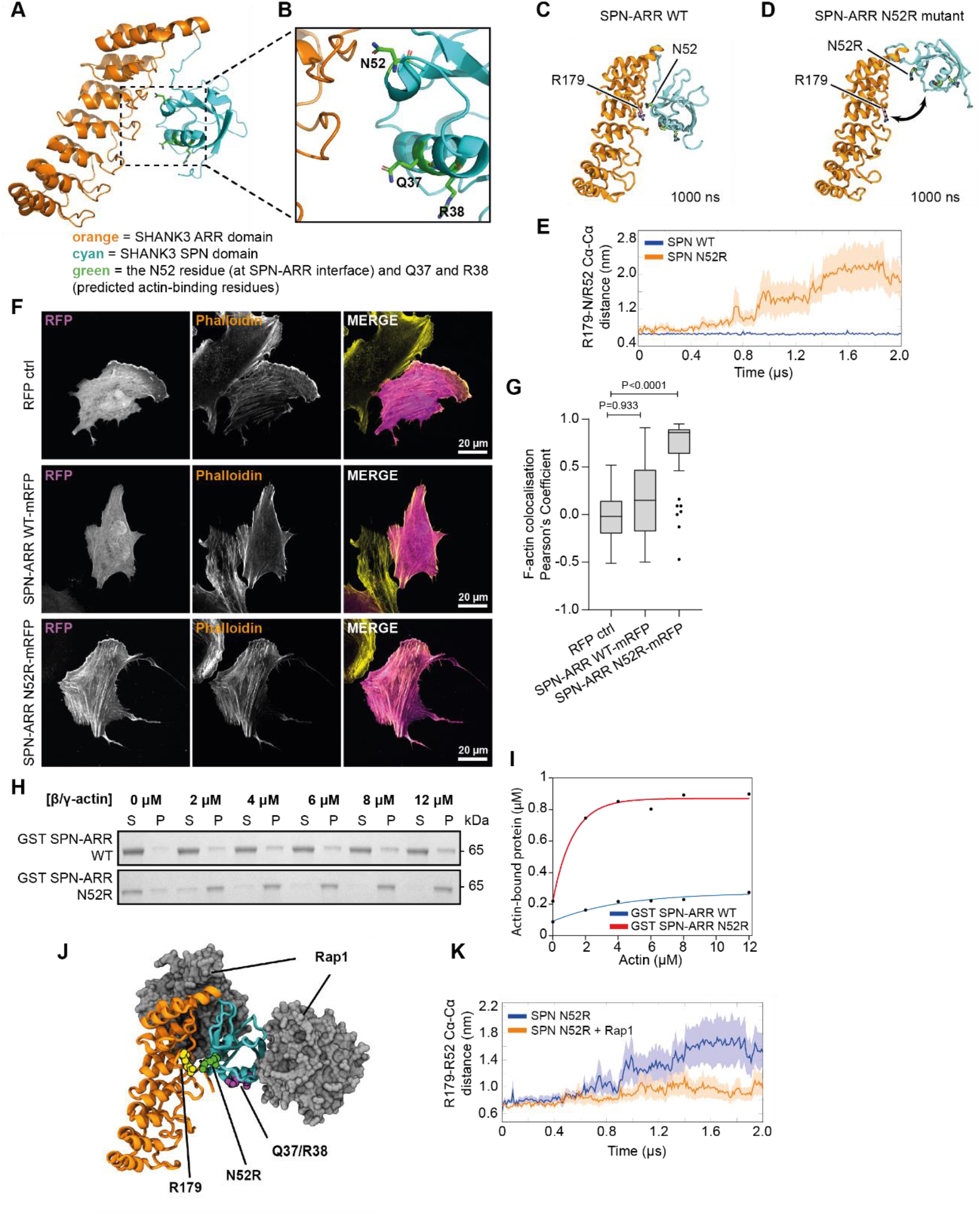
Mutation of the SHANK3 SPN-ARR interface induces an open conformation and promotes actin binding. **A, B,** Visualization of the SHANK3 SPN-ARR domains (5G4X, residues 1-348) using Pymol. Dashed area was expanded to illustrate the folding of the two domains and the close proximity of the SPN actin-binding site (residues Q37/R38) to the ARR-SPN interface. **C, D,** The structure of SPN-ARR WT (C) and N52R mutant (D) determined from atomistic MD simulations at 1000 ns. The snapshots are taken from the simulation systems S1 and S2 (see Table 1). **E,** Analysis of the distance between Cα atoms of residues N52 and R179 during the simulations. R179 was selected as it is located directly next to the N52 residue in both available X-ray structures (5G4X and 6KYK). The data are calculated from Systems S1 and S2 (Table 1). Standard errors are represented with shading. **F, G,** Analysis of localization between RFP and F-actin (phalloidin-647) in U2OS cells expressing either RFP control, SPN-ARR WT-mRFP or SPN-ARR N52R-mRFP and plated on fibronectin (3-4 h). Representative bottom plane confocal images (F) and Pearson’s correlation coefficient (G) quantified using coloc2 ImageJ plugin are shown. Two independent experiments. **H, I,** Analysis of GST-tagged SPN-ARR WT and GST SPN-ARR N52R binding to β/γ-actin filaments in a co-sedimentation assay (H) and quantification (I) analysed by SDS-PAGE and Coomassie Blue staining. The Exponential equation *f* = *y*_*o*_ + *a* * (1 − *exp*^−*b*\**x*^) was applied to fit experimental data. **J,** The structure of SPN-ARR N52R with two Rap1-GTP molecules taken from MD simulations (System S4, Table 1) at 1000 ns. **K,** Analysis of the distance between the Cα atoms of residues R179 and R52 as a function of simulation time. The data are calculated from Systems S3 (Table 1). Standard errors are represented with shading. All data are from three independent experiments unless otherwise indicated. Data represent mean ± s.d. (G). Number of cells: (G) 57 (RFP ctrl), 52 (SPN-ARR WT-mRFP) and 53 (SPN-ARR N52R-mRFP). Statistical analysis: (G) Kruskal-Wallis non-parametric test and Dunn’s multiple comparisons post hoc test. S, supernatant fraction; P, pellet fraction.

Based on the SPN-ARR structure ^14^, we predicted that mutating N52 (personal communication, Prof. Igor Barsukov, University of Liverpool, UK), a key residue at the SPN-ARR interface, may destabilize the closed conformation (Fig. 4A-B) and actin binding. Molecular dynamics simulations of the SPN-ARR WT and N52R mutant indicated that this mutation would trigger a conformational change in the molecule, exposing the actin-binding site (Fig. 4C-E). Based on the available structural data ^14,41^, we generated atomistic *in silico* models of the SPN-ARR region and modelled SPN-ARR WT (System S1 in Table 1 in the methods) and N52R mutant (System S2 in Table 1). Multiple independent 2 μs simulations of this model revealed dissociation and opening of the initially closed SPN-ARR interface in the N52R mutant (Fig. 4C-E, Supplementary Video 1) whereas the WT retained a closed conformation. Corroborating these findings, we calculated the affinity of SPN-ARR binding to be ~ ΔG_N52R_ = 21 kJ/mol lower with the N52R mutant compared to the WT (Extended Data Fig. 4D) (data based on free energy calculations; Systems S5 and S6 in Table 1). It is likely that the charge repulsion between R52 (SPN domain) and R179 (in the ARR domain) plays a role in the decreased stability of the interface in case of the N52R mutant, as no other differences were observed between WT and the N52R mutant during these simulations. These *in silico* data, indicating fold opening, were also supported by experimental data. SPN-ARR N52R-mRFP protein, but not SPN-ARR WT-mRFP, efficiently pulled down GFP SPN in cells (Extended Data Fig. 4E-F), indicating that the N52R point mutation exposes the ARR domain for subsequent binding to GFP SPN.

**Table 1.** Description of simulated systems.

| *System* | *Protein components*<br>*(mutation, residue range)* | *No. of water molecules*<br>*($K^+$, $Cl^-$)* | *No. of replicas x duration (ns)* | *PDB id.* |
| --- | --- | --- | --- | --- |
| *S1* | SHANK3 SPN-ARR (wt, 2–347) | 59354 (171, 169) | 4 x 2000 | 5G4X |
| *S2* | SHANK3 SPN-ARR (N52R, 2–347) | 59386 (170, 169) | 4 x 2000 | 5G4X |
| *S3* | SHANK3 SPN-ARR (N52R, 5–363) | 59330 (169, 173) | 4 x 2000 | 6KYK |
| *S4* | SHANK3 SPN-ARR (N52R, 5–363), 2 x Rap1A (wt, 1–166), 2x GNP | 65065 (193,185) | 8 x 2000 | 6KYK |
| *S5* | SHANK3 SPN-ARR (wt, 2–347) | 61674 (170, 168) | 7 x 300 | 5G4X |
| *S6* | SHANK3 SPN-ARR (N52R, 2–347) | 61672 (169, 168) | 7 x 300 | 5G4X |

Importantly, whereas WT SPN-ARR-mRFP displayed a diffuse cytoplasmic localization when expressed in U2OS cells, the SPN-ARR N52R-mRFP mutant displayed striking localization to F-actin rich structures (Fig. 4F-G). Moreover, recombinant SPN-ARR N52R co-sedimented efficiently with β/γ-actin filaments, whereas SPN-ARR WT displayed only very weak binding to actin filaments in this assay (Fig. 4H-I, Extended Data Fig. 4G). In addition, SPN-ARR N52R-mRFP pulled down β-actin from cell lysates as effectively as the positive actin-binding control mRuby LifeAct (Extended Data Fig. 4H), whereas β-actin was largely absent from WT SPN-ARR-mRFP pulldowns.

Recent structural data of SHANK3 N-terminus indicate that in addition to the canonical Rap1 binding site on the SPN domain, there is a second unconventional Rap1 binding site formed by SPN and ARR domains together ^41^. Therefore, we included Rap1 in the simulations and found that Rap1 inhibited SPN-ARR N52R opening (Fig. 4J-K, Supplementary Video 2). Imaging supported these data. Co-expression of active GFP Rap1 (Q63E) with SPN-ARR N52R-mRFP significantly reduced actin colocalization compared to cells co-transfected with GFP alone (Extended Data Fig. 4I-J). These data indicate that the SPN‒actin interaction is regulated dynamically by opening of the SPN-ARR fold, and that Rap1 inhibits SHANK3-actin interaction via two mechanisms: by controlling the opening of the SPN-ARR interface and by competing with F-actin binding to the SPN domain. We hypothesize that in cells, there is a physiological signal that triggers the opening of the SPN-ARR fold, but the nature of that signal remains to be investigated.

### The open SPN-ARR fold triggers full-length SHANK3 recruitment to actin filaments

To investigate the relevance of the SPN-ARR fold opening for SHANK3, we introduced the N52R point mutation into full-length GFP SHANK3. GFP SHANK3 N52R strongly localized to actin rich structures, such as stress fibers (Fig. 5A). This was markedly distinct from GFP SHANK3 WT and GFP SHANK3 Q37A/R38A, which localized to the plasma membrane and throughout the cytoplasm without clear accumulation to actin stress fibers (Fig. 5A) Within stress fibers, full-length GFP SHANK3 N52R displayed a periodic localization pattern, interspersed with non-muscle myosin IIA staining (Fig. 5B-C). Thus, also in the context of full-length SHANK3, opening of the SPN-ARR interface (N52R mutation) activates the actin-binding function that leads to accumulation of SHANK3 to actin filament rich structures in cells.

**Figure 5.**
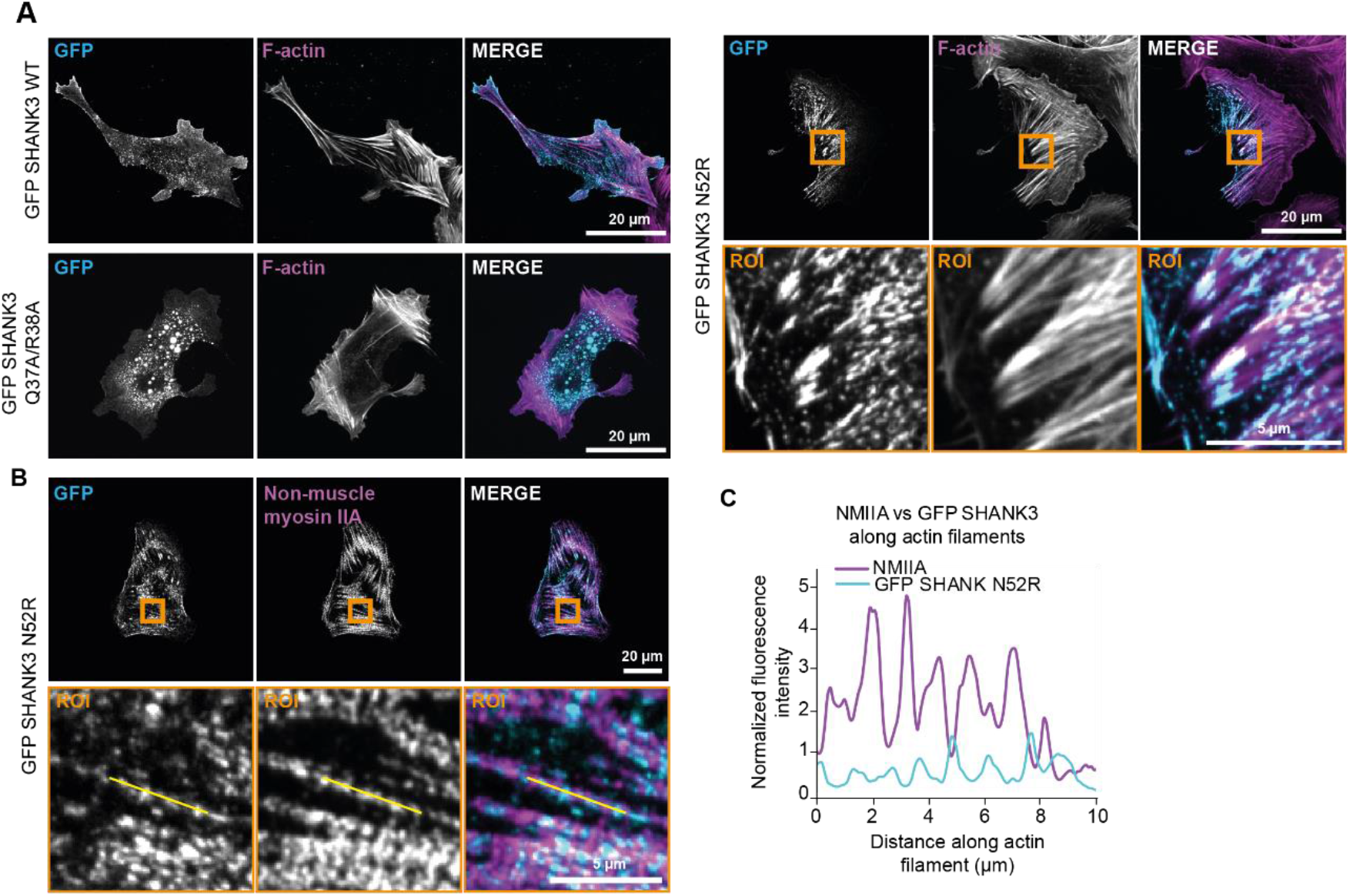
The SHANK3 N52R mutant localizes to actin stress fibers. **A,** Representative bottom plane confocal images of U2OS cells expressing either GFP control, GFP SHANK3 WT, Q37A/R38A or N52R stained with attophalloidin-647 to visualize F-actin. Cells were plated on fibronectin (3-4 h) before fixation. **B, C,** Analysis of the distribution of GFP SHANK3 N52R and endogenous NMIIA (non-muscle myosin IIA) along stress fibers in U2OS cells plated on fibronectin (3-4 h). Representative bottom plane confocal images (B) and a representative line scan along an actin stress fiber (C) are shown. Orange squares highlight ROI that are magnified.

### SHANK3-actin interaction modulates dendritic spine development

SHANK3 localizes to actin-rich dendritic spines where it acts as a major scaffolding molecule for actin regulatory proteins ^29,47^. *Shank3*-deficient mice have autism-like symptoms that can be rescued by restoration of *Shank3* in adult animals ^48^ or by targeting actin regulators ^30^. To explore if the SHANK3 SPN domain has a functional role in the development of dendritic spines, we expressed GFP SHANK3 WT and GFP SPN WT in primary hippocampal neurons isolated from WT rats. In mature neurons, spine density was not significantly different in any of the conditions. However, consistent with previous reports, the exogenous expression of GFP SHANK3 WT promoted the incidence of high spine density (Extended Data Fig. 5A) ^47^. In contrast, the expression of the SPN domain alone resulted in more neurons with medium or low spine density (Fig. 6A) and mature GFP SPN WT-expressing neurons exhibited a significantly lower spine head diameter to neck length ratio compared to GFP SHANK3 neurons (Fig. 6B-C). These data indicate that expression of the SPN domain alone has dominant negative effects on spine density and morphology and full-length SHANK3 or longer SHANK3 fragments are required for supporting normal spine development.

**Figure 6.**
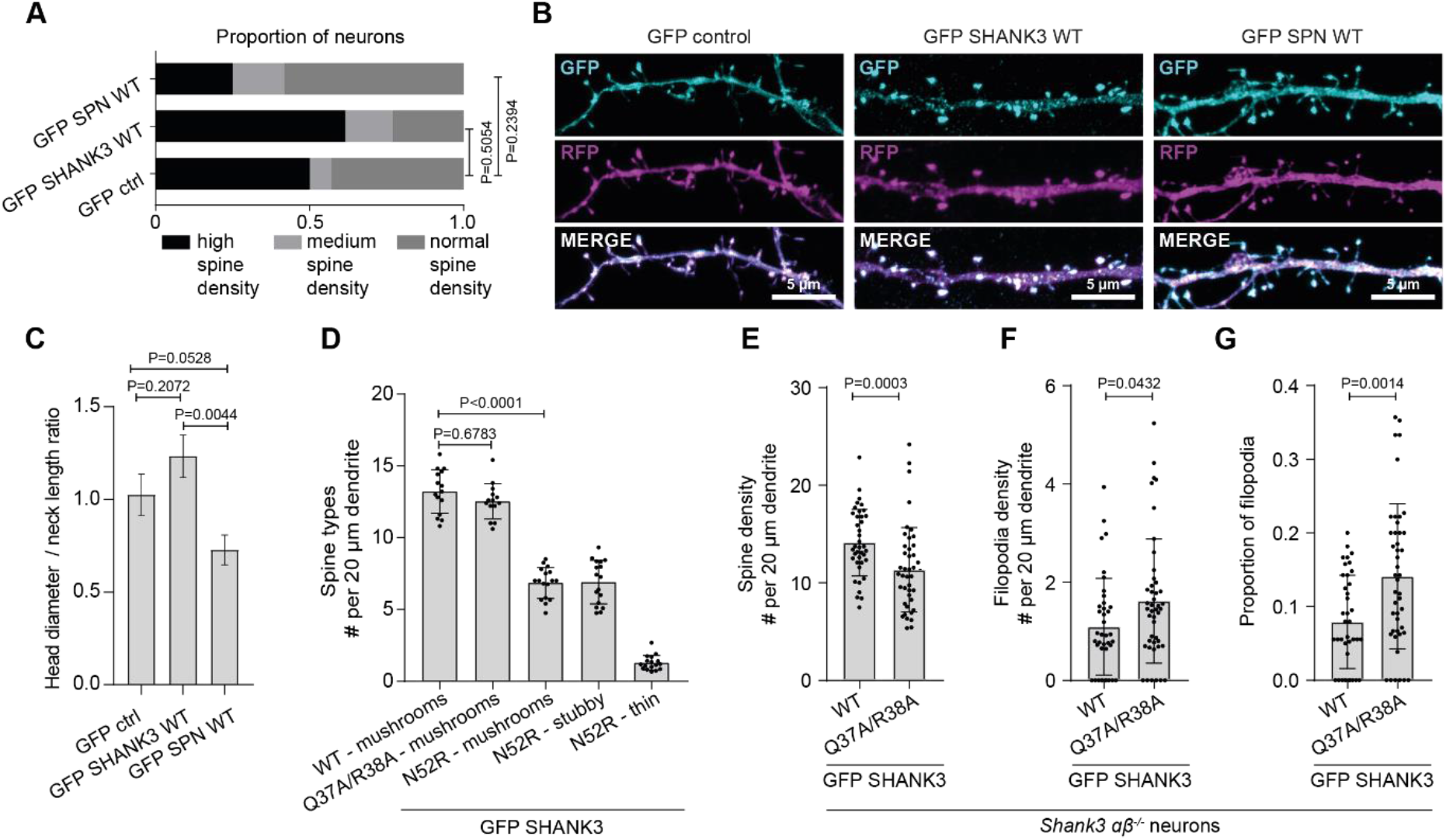
SHANK3-actin interaction regulates spine morphology and number. **A,** Quantification of spine density of WT primary rat hippocampal neurons fixed at DIV16-18. **B, C,** Representative maximum intensity projection confocal images (B) of WT primary rat hippocampal neurons fixed at DIV16-18 co-expressing RFP and GFP control, GFP SHANK3 WT or GFP SPN and (C) quantification of spine head diameter to neck length ratio. **D,** Analysis of WT primary rat hippocampal neurons expressing GFP SHANK3 WT, Q37A/R38A or N52R fixed at DIV16. Number of different spine types per 20 μm dendrite are shown. **E, F, G,** Analysis of spine development and filopodia formation in primary *Shank3αβ*^*−/−*^ mouse hippocampal neurons fixed at DIV14 expressing GFP SHANK3 WT or Q37A/R38A. Quantification of spine density (E), filopodia density (F) and proportion of filopodia (G) are shown. (A) Data represent the proportion of neurons in each spine density category. (C-G) Data represent mean ± s.d.; (A) n = 14 (GFP ctrl), 13 (GFP SHANK3 WT) and 25 (GFP SPN WT) neurons; (C) 14 neurons, 154 spines (GFP ctrl), 16 neurons, 223 spines (GFP SHANK3 WT) and 7 neurons, 104 spines (GFP SPN WT); (D) number of branches: 45 from 15 neurons; (E, F, G) Number of secondary dendrites: 39 (WT) and 45 (Q37A/R38A). Statistical analysis: (A) Chi-Square. (C) one-way ANOVA. (D) Kruskal-Wallis non-parametric test and Dunn’s multiple comparisons post hoc test. (E,F, G) Mann-Whitney two-tailed T-test.

To investigate whether direct binding to actin is required for the functionality of full-length SHANK3, we expressed GFP SHANK3 WT and the actin-binding mutants GFP SHANK3 Q37A/R38A and GFP SHANK3 N52R first in primary WT rat neurons expressing endogenous SHANK3 (Extended Data Fig. 5B). Spine density remained unchanged in all of the conditions tested (Extended Data Fig. 5C). However, GFP SHANK3 N52R-expressing neurons showed a striking 50 % decrease in the number of mushroom-shaped spines and a large proportion (40 %) of spines had a stubby morphology and appeared stretched on the dendritic shaft (Extended Data Fig. 5B, Fig. 6D). Mature mushrooms were the major spine type in GFP SHANK3 WT- and Q37A/R38A-expressing neurons, and the proportion of other spine types was negligible (therefore numbers not included in Fig. 6D). Despite their abnormal morphology, the spines of GFP SHANK3 N52R-expressing neurons were positive for vesicular glutamate transporter (vGlut) (Extended Data Fig. 5D). Thus, expression of GFP SHANK3 N52R did not affect the post-synaptic vGlut expression, but rather increased the prevalence of malformed stubby and thin spines at the expense of mushroom-shaped spines (Extended Data Fig. 5D, Fig. 6D). As the morphology of dendritic spines is believed to be largely determined by their actin cytoskeleton ^2,29,49^, these data indicate that the enhanced actin binding activity of SHANK3 N52R interferes with proper actin network formation in maturing dendritic spines. Expression of the actin-binding deficient GFP SHANK3 Q37A/R38A did not differ significantly from WT SHANK3 expressing rat neurons. We hypothesized that this may be due to the presence of endogenous SHANK3, given that SHANK3 homo-oligomerizes in the PSD ^50,51^. To overcome potential compensation by endogenous SHANK3, we expressed GFP SHANK3 WT and SHANK3 Q37A/R38A in neurons isolated from *Shank3αβ*^*−/−*^ mice that lack both the long α- and the shorter β-isoforms of *Shank3* ^52,53^. Neurons re-expressing GFP SHANK3 WT exhibited round spine heads, in keeping with earlier observations ^38,46,47^. In contrast, GFP SHANK3 Q37A/R38A-expressing neurons had lower spine density (Fig. 6E) and significantly higher number of filopodia compared to the GFP SHANK3 WT-expressing cells (Fig. 6F-G) indicative of a developmental delay. Taken together, these data suggest that direct SHANK3-actin interaction is required for normal SHANK3 function in neurons and that the enhanced actin binding of the N52R mutant interferes with maturation of dendritic spines even in the presence of endogenous WT SHANK3.

### SHANK3 actin-binding mutants are functionally defective in a zebrafish model of ASD

SHANK3 is well conserved in different species, and the zebrafish ortholog of human *SHANK3*, which exists in two copies (*shank3a* and *shank3b*), shares 55-68 % overall sequence homology with human *SHANK3*. Moreover, the sequence identity has been reported to be close to 100 % in many protein encoding regions ^54^. Transient morpholino-mediated knockdown of *shank3a* and *shank3b* expression or CRISPR/Cas9-mediated deletion of *shank3b* in zebrafish result in neurodevelopmental delay, including smaller brain, body and eye size, reduced eye pigmentation, as well as autism-like behavior such as repetitive swimming patterns, reduced locomotor activity and social interaction ^11,54,55^. Therefore, we employed zebrafish embryos to address whether SHANK3-actin interaction plays a role in early neurodevelopment. Knockdown of *shank3b* with morpholinos, significantly reduced eye pigmentation (Fig. 7A-B) consistent with previous reports ^55^. Introduction *of in vitro* transcribed GFP SHANK3 WT mRNA significantly rescued eye pigmentation whereas GFP SHANK3 N52R mRNA failed to rescue the phenotype (Fig. 7B).

**Figure 7.**
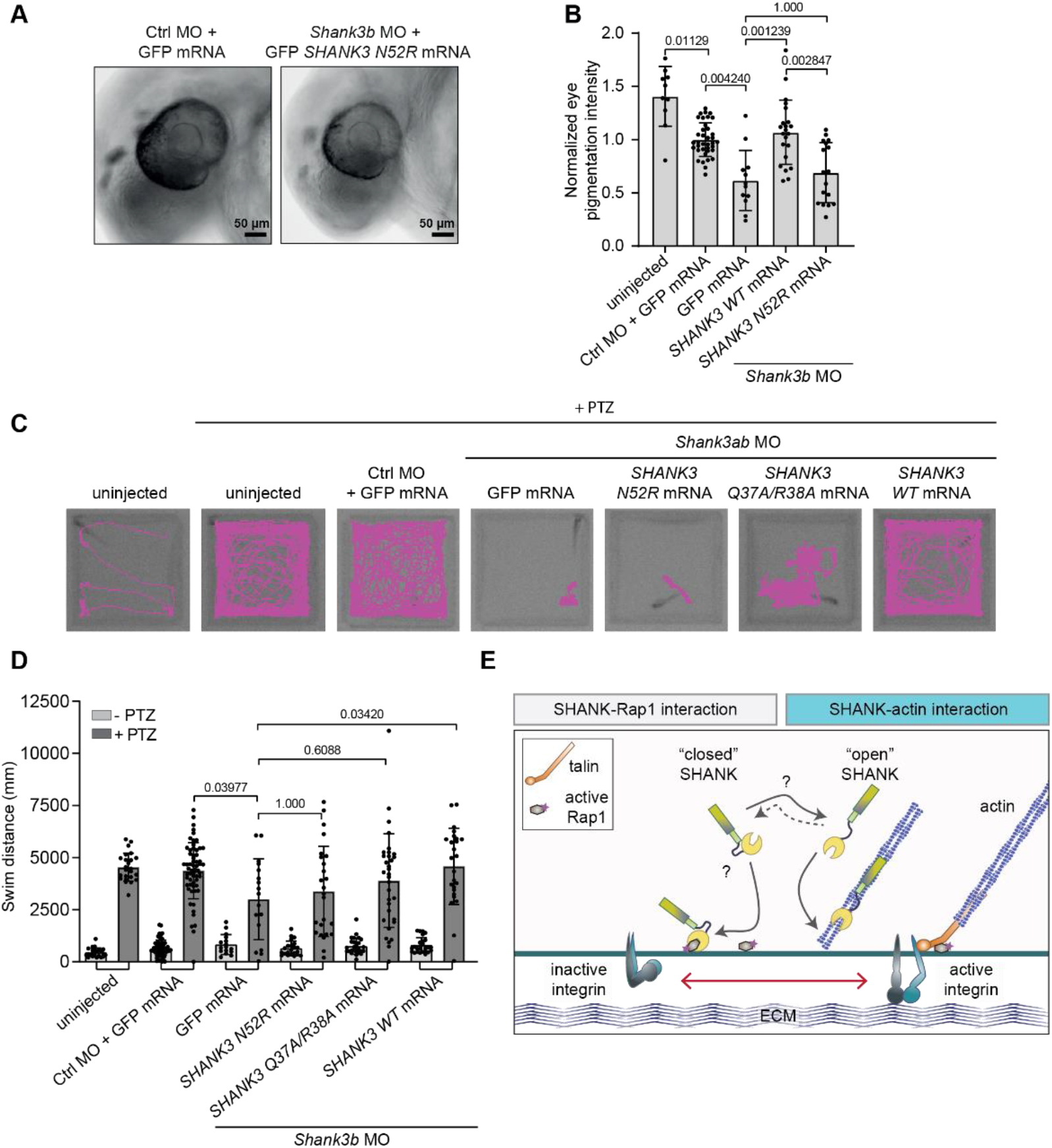
Dynamic SHANK3-actin binding is necessary for rescue of autism-linked phenotypes *in vivo.* **A, B,** Analysis of the eye pigmentation phenotype in zebrafish embryos microinjected with a *shank3b*-targeting morpholino (MO) and rescued with *in vitro* transcribed *SHANK3* mRNA co-injections. Pigmentation of the eye was analysed at 30 hpf. Images of the head of zebrafish embryos (A) and quantification of the pigmentation of the eye (B) are shown. **C, D,** Analysis of the motility of zebrafish embryos microinjected with *shank3a and b*-targeting morpholinos and rescued with *SHANK3* mRNA co-injections. Microinjected zebrafish embryos were dechorionated, placed in 96-well plates and subjected to motility analysis using the DanioVision instrument. After the first round of imaging, 20 mM pentylenetetrazole (PTZ) was added and samples analysed again. Zebrafish embryos were imaged at high-speed 30 fps and tracked automatically using Ethovision XT software. Recorded tracks of zebrafish embryo movement (C) are displayed in magenta and overlaid on the image of the 96-well plate. The total swimming distance (mm) of zebrafish embryos (D) is also shown. **E,** Summary figure. A SHANK3 conformational switch regulates SHANK3 actin and Rap1 binding and integrin activation. Number of embryos: (B) Control MO + GFP (37), *shank3b* MO + GFP (12), *shank3b* MO + N52R (17), *shank3a+b* MO + WT (22), uninjected (10). (D) (unstimulated/PTZ stimulated), Control MO + GFP (58 / 60), *shank3a+*b MO + GFP (16 / 17), *shank3a+b* MO + N52R (22 / 25), *shank3a+b* MO + QRAA (28 / 33), *shank3a+b* MO + WT (25 / 25), uninjected (24 / 24). Data are mean ± s.d. Statistical analysis: (B) non-parametric Kruskal-Wallis test and Dunn’s post-hoc test. (D) Rout’s outlier detection algorithm (Q=0.5%) followed by non-parametric Kruskal-Wallis test and Dunn’s post-hoc test.

Next, we analyzed zebrafish embryos in a motility assay (Fig. 7C-D). As mRNA rescue works most efficiently in early time points, we used 2 days post-fertilization embryos and utilized pentylenetetrazole (PTZ) to induce zebrafish embryo motility ^56^. In this assay, *shank3* knockdown resulted in reduced swim distance (Fig. 7D). Introduction of WT rat GFP SHANK3 mRNA rescued the effects on swim distance, but both GFP SHANK3 mRNAs carrying N52R or Q37A/R38A mutations failed to rescue this phenotype (Fig. 7D).These results suggest that mutations that either impair or enhance the actin-binding function of SHANK3 have dramatic loss-of-function effects on ASD-linked phenotypes *in vivo*.

## Discussion

Here, we reveal a cellular role for direct SHANK3 binding to actin, driven by a SHANK3 conformational switch that is inhibited by Rap1. We provide evidence for a dynamic crosstalk that has wide-ranging effects in governing actin-network architecture and integrin activity in cancer cells, dendritic spine morphology in neurons and functional implications for neuronal development in zebrafish embryos. SHANK3 binds active Rap1 and inhibits integrin activation ^14^. Dynamic regulation of the N-terminal SPN-ARR conformation by active Rap1 and other, yet unknown signals, enables SHANK3 coordinated crosstalk between integrin activity regulation and the actin cytoskeleton. A plausible scenario would be that when active Rap1 is highly abundant, SHANK3 binds to Rap1 sequestering it from the integrin activating Rap1-talin axis and the ‘closed’ SHANK3 conformation becomes stabilized. However, in areas of active actin polymerization and presence of actin filaments, ‘open’ SHANK3 favors actin binding and Rap1 is released promoting integrin activation. Thus, SHANK3 may play a key role in ensuring that Rap1-mediated integrin activation is restricted to actin-rich regions of the cell (Fig. 7E).

Understanding how the passage of information from adhesions to the actin cytoskeleton and back is mediated in a dynamic cell requires detailed understanding of the players involved. The Rap1 GTPase promotes activation of integrins ^32,57,58^, and integrin-mediated cell adhesion activates sequentially Rac and RhoA GTPases to induce actin polymerization, cell spreading and generation of stress fibers ^59,60^. On the other hand, actin and actin-binding proteins, such as talin, support integrin activity, receptor clustering and adhesion maturation ^61,62^. Therefore, coordination of integrin function and actin dynamics is expected to play a central role in the regulation of cell morphology and dynamics. However, very little is known about actin/adhesion crosstalk, especially in the context of specific adhesion types and actin-rich cell processes. Here, we report that SHANK3, which thus far has been considered as a scaffold for actin-binding proteins, interacts directly with F-actin through its N-terminal SPN domain. This SHANK3-actin interaction is necessary for the regulation of actin in dendritic spines. Our data indicate that SHANK3 may represent one important node connecting the dynamic regulation of the actin cytoskeleton with Rap1-mediated integrin activity.

SHANK3 expression promotes actin polymerization and increases F-actin levels in dendritic spine. This has been largely attributed to the ability of SHANK3 to recruit different actin regulators, such as Abl1 and the WAVE complex ^19^, β-PIX ^23^, cortactin ^26,27^ and IRSp53 ^25^, to the PSD. The interaction of the SHANK3 SPN domain with actin did not seem to modulate actin dynamics directly. Thus, it is plausible that the main function of SHANK3-actin interaction is to co-ordinate integrin activation with the actin cytoskeleton, and to recruit SHANK3-associated actin regulators to actin filaments. Lack of SHANK3 leads to poor development of dendritic spines ^29,30,47^. In line with these observations, the autistic phenotype caused by lack of *Shank3* in mice can be rescued by targeting actin regulators, namely inhibiting cofilin or activating Rac1 ^30^. S*hank3* depletion with morpholinos or CRISPR/Cas9 genome editing results in autism-like phenotypes and neuronal defects in zebrafish embryos and adult zebrafish 11,54,55,63, recapitulating the *SHANK3* loss-of-function phenotypes in mice and man. We find that augmented SHANK3-actin binding (N52R open-conformation mutant) and defective actin-binding (Q37A/R38A mutant) fail to rescue SHANK3 loss *in vivo* in the zebrafish. In addition, SHANK3 N52R shows dominant-negative activity over endogenous SHANK3 by suppressing normal dendrite morphology in neurons *in vitro* and by further deteriorating the motility of the *shank3*-depleted zebrafish embryos. These data indicate that the direct interaction between the SHANK3 SPN domain and F-actin is a key mechanism in SHANK3-mediated actin regulation.

Here, we establish a regulatory function for the SHANK3 SPN-ARR fold in controlling SHANK3 recruitment to actin filaments in cells. The N-terminal SPN-ARR is folded in a closed conformation *in vitro* ^14,46^. This fold has been shown to inhibit the binding of SHANK3-interacting proteins SHARPIN and α-fodrin to the ARR domain ^46^ and we find that the closed SPN-ARR does not interact with actin. Furthermore, molecular simulations indicate that this closed confirmation is stabilized by Rap1 binding. Collectively, these data suggest that the SPN-ARR fold opening and actin binding are dynamically regulated by Rap1 activity. In addition, we hypothesize that in cells, a physiological signal, such as post-translational modification, co-factor recruitment or interaction with membrane lipids, triggers opening of the fold and presumably spatially controls SHANK3-actin interaction. For example, the interaction between SHANK3 and Abl1 is regulated by phosphorylation at S685, a residue in the PP-domain, and an ASD-linked patient mutation S685I interferes with this phosphorylation abolishing interaction with Abl1 and decreasing downstream actin polymerization ^19^. However, the identity of the signal(s) regulating the SPN-ARR fold opening remains to be determined.

Other proteins have previously been shown to regulate integrin activity and bind actin. These include well-established integrin activators talin and kindlin ^61^. These proteins are, however, activators not inhibitors of integrins and actin binding does not directly affect their integrin activation properties. SHANK3 is unique in that its ability to inhibit integrin activity is coupled directly to actin binding. This would enable it to locally co-ordinate Rap1-signaling and integrin activity in response to changes in actin polymerization and vice versa (Fig. 7E). Given the undisputed biological importance of SHANK3 function in human health, SHANK3 is a prime candidate to fine-tune numerous physiological processes from neuronal actin regulation to cell migration in multiple other tissues. In this respect, dissection of the mechanisms regulating SHANK3 in physiology and pathology is a major challenge ahead of us.

## Acknowledgements

We thank the Cell Imaging and Cytometry and the Zebrafish Core facilities (Turku Bioscience Centre, University of Turku and Åbo Akademi University) and University of Turku Central Animal Laboratory all supported by Biocenter Finland for services in imaging and animal experiments. We thank P. Laasola, J. Siivonen and V. Pollari for technical help and H. Hamidi for editing of the manuscript and illustrations. I. Barsukov, J. Pouwels, P. Hotulainen and the Ivaska lab are acknowledged for important suggestions, feedback and stimulating discussions. This study has been supported by an ERC CoG grant 615258 (J.I.), the Academy of Finland (J.I., G.J. E.T.C., P.L.), the Finnish Cancer Foundation (J.I., P.L.), the Sigrid Juselius Foundation (J.I., G.J.) and by grants from Deutscher Akademischer Austauschdienst (DAAD; to F.H.-N.) and Deutsche Forschungsgemeinschaft (DFG; KR 1321/9-1; to H.-J.K.). S.I.S. has been supported by the Svenska kulturfonden, Finnish Cultural Foundation, University of Turku Foundation and Maud Kuistila Memorial Foundation. S.I.S. and M.M. have been supported by University of Turku Drug Research Doctoral Programme, J.L. by Turku Doctoral Programme of Molecular Medicine and A.I. by Turku Doctoral Programme of Molecular Life Sciences.

## Author contribution

Conceptualization, S.I.S., J.L., G.J., I.V., P.L., and J.I.; Methodology, S.I.S., E.K., M.M., J.V., G.J., J.L., F.H-N., I.P. and J.I.; Formal Analysis, S.I.S., E.K., L.A., J.V., J.L., P.H., F.H., I.P. and J.I.; Investigation, S.I.S., E.K., J.L., J.V. P.H., F.H-N., G.J., M.M., I.P. and J.I.; Writing – Original Draft, S.I.S. and J.I.; Writing – Review and Editing, S.I.S., H.J.K., P.L., J.V., I.V., E.T.C. and J.I.; Supervision, H.J.K., I.V., E.T.C., P.L., and J.I.; Funding Acquisition, H.J.K., P. L., E.T.C., and J.I.

## Data Availability

The data that support the findings of this study are available within the paper [and its supplementary information files] or from the corresponding author upon reasonable request.

## METHODS

### Cell lines and cell culture

CHO (Chinese hamster ovary) cells were grown in α-MEM medium (Sigma-Aldrich) supplemented with 5 % fetal bovine serum (FBS, Gibco), 2 mM L-glutamine (Sigma-Aldrich) and 1 % (vol/vol) penicillin/streptomycin (pen/strep, Sigma-Aldrich). HEK293 (human embryonic kidney) and U2OS (human bone osteosarcoma) cells were maintained in Dulbecco’s modified Eagle’s medium (DMEM, Sigma-Aldrich) supplemented with 10 % FBS, 2 mM glutamine and 1 % pen/strep. All cell lines were regularly checked for mycoplasma contamination. All cell lines were obtained from ATCC, except for U2OS, which were provided by DSMZ (Leibniz Institute DSMZ-German Collection of Microorganisms and Cell Cultures, Braunschweig DE, ACC 785).

### Mice and rats

Sprague-Dawley rats and Wistar Unilever outbred rats (strain HsdCpb:WU, Envigo, Horst, The Netherlands) were used for isolation of primary hippocampal neurons. *Shank3 αβ*-deficient mice were provided by Tobias Boeckers (Univ. of Ulm, Germany) (Schmeisser et al., 2012).

Timed, pregnant animals were housed in individual cages, with access to food and water ad libitum. All animal experiments were approved by, and conducted in accordance with, the Turku Central Animal Laboratory regulations and followed national guidelines for Finnish animal welfare, or regulations of the Animal Welfare Committee of the University Medical Center (Hamburg, Germany) under permission number Org766.

### Isolation and culture of primary hippocampal neurons

Newborn Sprague-Dawley rats were decapitated and their hippocampus was placed into dissection media (1 M Na_2_SO_4_, 0.5 M K_2_SO_4_, 1 M MgCl_2_, 100 mM CaCl_2_, 1 M Hepes (pH 7.4), 2.5 M Glucose, 0.5 % Phenol Red). Meninges were removed and hippocampal pieces were collected into dissection media containing 10 % KyMg, followed by washing. Hippocampal tissue was then incubated with 10 U/ml papain (#3119, Worthington) for 15 min at 37°C, repeated two times. Papain was inactivated by incubation with 10 mg/ml trypsin inhibitor (Sigma, T9128) for 2 × 5 min at 37°C. Hippocampal tissue was then homogenized by gentle pipetting. Cultures were plated on 0.1 mg/ml poly-D-lysine-coated glass coverslips and maintained in Neurobasal-A medium (Thermo Fisher Scientific) supplemented with 2 mM glutamine, 50 U/ml penicillin, 50 μM streptomycin and B27 Neuronal supplement (Gibco, Thermo Fisher Scientific).

Pregnant Wistar rats (Envigo; 4-5 months old) were sacrificed on day E18 of pregnancy using CO_2_ anaesthesia, followed by decapitation. Neurons were prepared from all embryos present, regardless of gender (14-16 embryos). The hippocampal tissue was dissected, and hippocampal neurons were extracted by enzymatic digestion with trypsin, followed by mechanical dissociation. Cells were grown in Neurobasal medium supplemented with 2 % B27 Neuronal Supplement, 1 % GlutaMAX and 1 % pen/strep (Gibco, Thermo Fisher Scientific) on 0.1 mg/ml poly-L-lysine-coated glass coverslips. Neurons were transfected using the calcium phosphate method as described below. Neurons from *Shank3 αβ*-deficient mice were isolated and transfected in a similar manner, except that the pregnant mice were sacrificed at E17 and neurons were analysed at DIV 14 since *Shank3 αβ*^−/−^ neuron cultures are more fragile in vitro.

### Plasmids

The SHANK3-mRFP (pmRFP-N3, Clontech) was described earlier ^14^, and deletion constructs generated either by using appropriate restriction sites or PCR amplification of cDNA fragments prepared in pmRFP-N3 vectors ^64^. The tD-tomato-N1 vector was obtained from Clontech. The construct coding for a GFP-fusion of the SHANK3 SPN domain has been described previously ^14,46^. A construct coding for N-terminal GFP-tagged full-length rat SHANK3 in the pHAGE vector was obtained from Alex Shcheglovitov (Univ. of Utah, Salt Lake City) ^14,65^. Constitutively active human Rap1A (pEGFP-C3-Rap1Q63E, here referred to as GFP Rap1 Q63E) was a gift from B. Baum and S. Royale ^66^. Myo10-mCherry was a gift from S. Strömblad, kindlin-2-GFP from M. Parsons and GFP-talin-1 from B. Goult. mRuby-Lifeact was obtained from Addgene (#54560). pEGFP-C1 and mRFP-N1 were used as controls in this study.

Bacterial expression constructs coding for Hi_s6_/SUMO tagged fusion proteins of rat SHANK3 were generated in pET-SUMO (Thermo Scientific) as described ^14^. For expression as glutathione-S-transferase (GST) fusion proteins, parts of the rat SHANK3 cDNA were amplified by PCR with oligonucleotide primers carrying appropriate restriction sites. Amplified fragments were subcloned into pGEX-2T or pGEX4T2 vectors (GE Healthcare) in frame with the GST coding sequence. Different point mutations were introduced into SHANK3 constructs by site-directed mutagenesis (Gene Universal) or by using mutagenic oligonucleotides and the Quick-Change II site-directed mutagenesis kit (Agilent) according to the manufacturer’s instructions. All modified plasmids were verified by sequencing before use.

### Transient transfections

Plasmids were transiently transfected into CHO, HEK293, and U2OS cells using Lipofectamine 3000 and P3000™ Enhancer Reagent (Thermo Fischer Scientific Inc, # L3000-015) according to manufacturer’s instructions. Cells were cultured for 24 h before they were re-plated (plating times indicated in figure legends) in subsequent experiments.

Primary neurons were either transfected at DIV16 with Lipofectamine™ 2000 Transfection Reagent (Thermo Fischer Scientific Inc, #11668019) according to manufacturer’s instructions, or using the calcium phosphate method at DIV7-9. For the latter, the complete Neurobasal medium was collected from wells one hour before transfection and replaced with pre-warmed transfection medium (MEM+GlutaMAX). Plasmid DNA was diluted in H_2_O and mixed with 2.5 M CaCl_2_. An equal amount of 2X Hepes buffered salt solution (HBS) was added drop-wise to the reaction tube under continuous mixing. The reaction was incubated at room temperature (RT) for 30 min and then divided between the wells of the cell culture plate. After a 2 h incubation with the transfection mixture, the cells were washed seven times with 1×Hank’s Balanced Salt Solution (HBSS). After the final wash, the previously collected Neurobasal medium was added back to the cells. 2xHBS: NaCl 274 mM; KCl 10 mM; Na_2_HPO_4_ 1.4 mM; D-Glucose 15 mM; Hepes 42 mM; adjusted to pH 7.05 with NaOH.

### Antibodies and reagents

Antibodies were used to detect proteins in western blots (WB) and immunofluorescence (IF) microscopy. Mouse antibodies used in the study were raised against β-actin (Sigma-Aldrich, Clone AC-15, # A1978, 1:1000 for WB), GFP (Abcam, #ab1218, 1:1000 for WB and Covance, #MMS-118P-500, 1:3000 for WB) and vinculin (Sigma-Aldrich, #V9131, 1:500 for IF). Rabbit antibodies were anti-GFP (Abcam, #ab69507, 1:1000 for WB), anti-GST (Cell Signalling Technology, #91G1), anti-non-muscle myosin heavy chain II (BioLegend, clone Poly19098, #909801, 1:1000 for IF), anti-RFP (Invitrogen, #R10367, 1:1000 for WB and Chromotek #5F8, 1:1000 for WB), anti-phospho myosin light chain (Cell Signaling Technology, #3674, 1:100 for IF) and anti-vesicular glutamate transporter (vGlut1) (Synaptic Systems; 1:2000 for IF). Alexa Fluor 488, 555 and 568-conjugated secondary antibodies (Invitrogen) against mouse and rabbit were used 1:300 for IF. IrDye 680 and IrDye 800 (LI-COR, 1:5000) against mouse and rabbit or HRP-labeled goat secondary antibodies (Jackson ImmunoResearch, 1:2500) were used in WB. F-actin was visualized using directly conjugated actin dyes Atto-Phalloidin-647 (Sigma, #65906, 1:500 for IF), Atto-Phalloidin-740 (Sigma, #07373, 1:75 for IF), Phalloidin Alexa Fluor 488 (Invitrogen, #A12379, 1:200 for IF), Phalloidin Alexa Fluor 647 (Invitrogen, #A22287, 1:200 for IF) and SiR-actin-647 (Spirochrome, #SC001, 1:5000 for IF).

### Immunofluorescence, confocal imaging and image analysis

For the immunofluorescence experiments with cell lines, 35 mm #1.5 glass-bottom dishes (Cellvis, #D35-14-1.5-N) were coated with bovine plasma fibronectin (Merck-Millipore, #341631, diluted to 10 μg/ml in PBS) overnight at +4 °C. Cells were plated on dishes in the appropriate medium for the indicated times. Cells were then fixed and permeabilized simultaneously by adding 16 % (wt/vol) paraformaldehyde and 10 % (vol/vol) Triton-X directly into the media at a final concentration of 4 % PFA and 0.1-0.25 % (vol/vol) Triton-X for 5-10 min, after which samples were washed with PBS and quenched with 1 M glycine in PBS for 25 min. Samples were incubated with primary antibodies (30 min at RT), followed by washes and incubation with fluorescently-conjugated secondary antibodies for 30 min at RT. Unless otherwise stated, the bottom plane was imaged with a Marianas spinning disk confocal microscope (3iw1) equipped with a CSU-W1 scanner (Yokogawa) and Hamamatsu sCMOS Orca Flash 4.0 camera (Hamamatsu Photonics K.K.) using a 63x/NA 1.4 oil, Plan-Apochromat, M27 with DIC III Prism objective. For images acquired using the structured illumination microscope (SIM), cells were plated on high tolerance glass-bottom dishes (MatTek Corporation, coverslip #1.7). Samples were fixed, permeabilized and stained as described above. Just before imaging, samples were washed three times in PBS and mounted in vectashield (Vector Laboratories). The SIM used was DeltaVision OMX v4 (GE Healthcare Life Sciences) fitted with a 60x Plan-Apochromat objective lens, 1.42 NA (immersion oil RI of 1.516) used in SIM illumination mode (five phases x three rotations). Emitted light was collected on a front-illuminated pco.edge sCMOS (pixel size 6.5 mm, readout speed 95 MHz; PCO AG) controlled by SoftWorx.

Primary neurons were grown on glass coverslips and fixed at indicated DIV with 4 % PFA followed by permeabilization with 0.1-0.5 % Triton-X and blocking with 10 % horse serum in PBS. Neuron samples were stained as described above and imaged either with an LSM880 Airyscan laser-scanning confocal microscope (Zeiss) with Airyscan detector using 63x/ 1.4 oil objective, or with a Leica TCS SP5 confocal microscope with 63x/1.4-0.60 HCX PL APO Lbd. Bl. oil objective.

Quantitative image analysis was performed with ImageJ. Colocalization analysis was done with ImageJ Coloc 2 plugin.

### Micropatterns

Crossbow-shaped micropatterns were created on acid-washed glass coverslips as previously described ^67,68^. The patterns were coated with 50 μg/ml fibronectin and 1:200 555-labelled BSA or 647-labelled fibrinogen before plating the cells. Cells were fixed 3-4 h after plating as described above.

### Analysis of dendritic spines

For dendritic spine head-and-neck ratio measurements, ImageJ’s line measurement tool was used on maximum intensity projection images and at least 16 spines were selected randomly from each cell. For neck length, a line was drawn and distance was measured from the base of the neck to the stem of the spine head. Head diameter was estimated by measuring the distance of a line between the two most distant points on the spine head. Head diameter/neck length ratios were calculated accordingly using Microsoft Excel. Spine density was analysed from Z-stacks using Neurolucida Explorer (MBF Bioscience, Williston, DC, USA) or with ImageJ. In *Shank3αβ*^−/−^ neurons, filopodia and other spine types were categorised manually on the basis of morphology of spines and whether they had a visible neck and a separate bulbous head (spine) or no apparent head at all (filopodia).

### Expression and purification of recombinant proteins

Competent E. coli BL21 bacteria were transformed with IPTG-inducible expression constructs having either GST- or SUMO-tag (also includes a His-tag) and grown in LB medium supplemented with selection antibiotics (ampicillin or kanamycin), at 37°C until an OD600 of 0.6-0.8. Protein production was induced by the addition of 0.1 mM IPTG overnight at 18 °C. The next day, the bacterial pellet was harvested by centrifugation for 20 min at 6000 g and then resuspended in cold lysis buffer (50 mM Tris, 150-300 mM NaCl, cOmplete™ protease inhibitor tablet (Roche, #5056489001) and 2 μl/ml DNAse (Sigma-Aldrich, #11284932001)). A small spoonful of lysozyme from chicken egg white (Sigma-Aldrich, #L6876-5G), 1 % Triton-X and 1x BugBuster (Merck Millipore, #70584-4) were added to lyse the bacteria for 30 min at 4 °C with gentle rotation. The lysate was cleared by centrifugation at 15000 g for 1 h at 4 °C. The cleared lysate was incubated with either Glutathione Sepharose® 4B (for GST-tagged proteins, GE Healthcare, #17-0756-01) or Protino Ni-TED resin (for SUMO-tagged proteins, Macherey-Nagel, #745200.5) for 1 h at 4 °C with rotation and then transferred to gravity columns (Talon® 2 ml Disposable Gravity Column, Clontech, #635606-CLI). The lysate was drained and the beads were washed five times with cold wash buffer (50 mM Tris, 150-300 mM NaCl). Elution buffers were made by adding 1 mM DTT (Sigma-Aldrich, #D0632-5G) and 0.1 % triton-X to the wash buffer. Furthermore, 30 mM reduced L-Glutathione (Sigma-Aldrich, #G4251-25G) or 250 mM imidazole was added to elute GST- or SUMO-tagged proteins, respectively. After addition of the eluting agent, the pH was adjusted to 7.0-7.4. Proteins were further dialyzed with Thermo Scientific Slide-A-Lyzer™ Dialysis Cassettes. Eluted and dialyzed proteins were analyzed with SDS PAGE gel electrophoresis and Coomassie Blue staining (InstantBlue Protein Stain, expedeon, #ISB1L).

### Co-sedimentation assays

Actin co-sedimentation assays were carried out essentially as described earlier ^69^. Briefly, different amounts of β/γ-actin were polymerized for 30-40 minutes at RT in the presence of G-buffer (5 mM Tris-HCl pH 7.5, 0.2 mM DTT, 0.2 mM CaCl_2_, 0.2 mM ATP) by addition of 5 mM MgCl_2_, 1 mM EGTA, 0.2 mM ATP, 1 mM DTT and NaCl at a final concentration of 100 mM. 1 μM of GST- or SUMO-tagged SPN/SPN-ARR WT or mutant variants in their respective buffers (for GST-tagged proteins: 50 mM Tris-HCl pH 8.0, 150 mM NaCl, 1 mM DTT and 0.1 % triton-X; for SUMO-tagged proteins: 50 mM Tris-HCl pH 8.0, 300 mM NaCl, 1 mM DTT and 0.1 % triton-X) were added to pre-polymerized actin samples and further incubated for 30 minutes. To sediment the polymerized actin filaments and bound proteins, the samples were subjected to ultracentrifugation for 30 minutes at 20 °C in a Beckman Optima MAX Ultracentrifuge at 60000 rpm for SUMO SPN-ARR WT, GST SPN WT and SPN Q37A/R38A and at 50000 rpm for GST SPN-ARR WT and SPN-ARR-N52R using a TLA100 rotor. Equal proportions of supernatants and pellets were run on 4-20 % gradient or 12 % SDS-polyacrylamide gels (Mini-PROTEAN TGX Precast Gels, Bio-Rad Laboratories Inc.), which were then stained with Coomassie Blue. The intensities of protein bands were quantified with ImageLab 6.0 program (Bio-Rad Laboratories Inc.), analyzed and plotted as actin-bound protein (μM, SPN protein in pellet) vs actin concentrations. Binding curves were fitted with 3 parameter exponential equation using SigmaPlot 11.0: *f* = *y*_*o*_ + *a* * (1 − *exp*^−*b*\**x*^), Binding curves were fitted with 3 parameter exponential equation using SigmaPlot 11.0: where *f* is actin-bound protein in μM, *y*_*o*_ is the protein in the pellet in the absence of actin, *a* the maximum bound protein, *x* actin concentration in μM and *b* is the fitting parameter. Actin concentration when half of the protein is bound was estimated from the equation: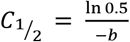.

To analyze the competition between actin and His-Rap1b (Cytoskeleton Inc, cat. no. RR02-A) biding to GST SPN some modifications were made to the assay. First, His-Rap1b was converted to active form by loading with a 10-fold excess of GMPPCP (non-hydrolyzable analogue of GTP, #M3509-25MG, Sigma-Aldrich) for 24 h at +4 °C in Exchange buffer (20 mM Tris-HCl pH 7.5, 150 mM NaCl, 10 mM MgCl_2_, 1 mM DTT, 5 % sucrose and 1 % dextran). After incubation, the buffer was changed to modified GST-buffer (50 mM Tris-HCl pH 8.0, 300 mM NaCl, 5 mM MgCl_2_, 5 % glycerol, 0.5 mM DTT and 0.1 % Triton-X) using Amicon buffer-exchange filters. Co-sedimentation assays were always performed with freshly made active His-Rap1b. 12 μM of β/γ-actin was polymerized for 1 hour at RT, followed by incubation with His-Rap1b and GST SPN, added sequentially, for approx. 50 min at RT. The final NaCl concentration in samples was always maintained at 100 mM. Then samples containing different combinations of actin, SPN and Rap1b proteins were sedimented for 30 minutes at 20 °C in a Beckman Optima MAX Ultracentrifuge at 60000 rpm in a TLA100 rotor. Equal proportions of carefully separated supernatants and pellets were run on 12 % SDS-polyacrylamide gels, which were processed as described above. The intensity values for SPN and Rap1b were corrected using values of similar-sized-bands from actin-alone and actin-SPN samples before further quantification, because of minor contaminants in the actin prep.

### β/γ-actin disassembly assay

The steady-state rate of β/γ-actin filament disassembly was measured using a modified protocol described for muscle actin (Kremneva et al., 2014). Samples of polymerized pyrene actin (4 μM) were mixed and incubated for 5 minutes with 1 or 2 μM GST-SPN and 0.8 μM cofilin-1 both diluted with G-buffer (5 mM Hepes pH8, 0.2 mM CaCl_2_, 0.2 mM ATP, 1 mM DTT), in the presence 0.8 μM cofilin-1 and in the absence of both. All protein mixtures were assembled in 1.5 ml Eppendorf tubes. The reaction was initiated by the addition of 6 μM vitamin D binding protein [DBP] (Human DBP, G8764, Sigma) directly in the fluorometric cuvettes. During the experiments, buffer conditions were constant: 20 mM HEPES pH 8, 100 mM KCl, 1 mM EGTA, 0.2 mM ATP. All measurements were carried out using the Agilent Cary Eclipse Fluorescence Spectrophotometer with BioMelt Bundle System (Agilent Technologies) with excitation at 365 nm (Ex. Slit = 5 nm) and emission at 407 nm (Em. Slit = 10 nm). Each measurement was carried out in triplicate.

### Co-immunoprecipitation

GFP-Trap® agarose, RFP-Trap® agarose and RFP-Trap® magnetic agarose (ChromoTek, #GTA-100, RTA-100 and RTMA-100) were used to pull down GFP- and RFP-tagged proteins from cell lysate. HEK293 and U2OS cells were transfected as described earlier, lysed in IP lysis buffer (40 mM HEPES-NaOH, 75 mM NaCl, 2 mM EDTA, 1% NP-40 and protease and phosphatase inhibitor tablets). Lysates were cleared by centrifugation and incubated with 30 μl of beads for 1 h at 4°C with rotation. The co-immunoprecipitated complexes were washed three times with the GFP IP wash buffer, resuspended in denaturating and reducing 4X Laemmli sample buffer and heated for 5 min at 95°C. GST- and SUMO-tagged recombinant proteins were bound to GSH sepharose and Macherey-Nagel Ni-Ted resin as described earlier and pull down assays were performed similarly to other co-immunoprecipitations, except for the IP wash buffer recipe which consisted of 20 mM Tris-Hcl (pH 7.5), 150 mM NaCl and 1 % NP-40. Samples were analyzed by SDS-PAGE followed by western blot.

### Western Blot and Coomassie Blue staining

Purified recombinant proteins and protein extracts prepared from harvested cells or immunoprecipitation experiments in reducing Laemmli Sample Buffer were run on 4–20 % Mini-PROTEAN® TGX™ Precast Protein Gels of different comb and well-sizes (Bio-Rad, #456-1093, #456-1094, #456-1095, #456-1095). For western blotting, gels were transferred to 0.2 μm nitrocellulose Trans-Blot Turbo Transfer Pack, mini or midi format (Bio-Rad, #170-4158, #170-4159). After transfer, membranes were blocked in 1:1 PBS and Thermo Scientific™ Pierce™ StartingBlock™ (ThermoFisher Scientific, #10108313). Primary antibodies were incubated overnight at +4 °C, and secondary antibodies for 1 h at RT, both in rotation or shaking. All antibody dilutions were done in the blocking buffer. Membranes were washed between antibody additions and before detection with Tris-buffered saline with Tween® 20 (TBST) and stored in PBS. Alternatively, samples were run on self-cast 10 % gels, blotted on nitrocellulose membranes using Wet Blot, blocked with and stained in 5 % milk in TBST and detected using WesternBright ECL Western Blotting detection kit (#K-12045-D20, Advansta). For Coomassie Blue staining, the gels were stained with Instant Blue (Biotop, #ISB1L) according to manufacturer’s instructions. The Odyssey (LI-COR) infrared scanner and Bio-Rad Chemidoc were used to image membranes and gels.

### Protein structure visualization and structure-based superimpositions

Pymol (The PyMOL Molecular Graphics System, Version 2.0 Schrödinger, LLC) together with the protein structure database (rcsb.org) were used to visualized different protein domains. Sequence alignment followed by structural superposition was carried out by using Pymol’s align-function. In cases of low sequence homology, Pymol’s cealign function was used instead. Pymol was used under professional license for academics.

### Multiple sequence alignment

MUSCLE multiple sequence alignment algorithm inside Geneious R8 (https://www.geneious.com) was used to align multiple protein sequences. Altogether, the Geneious software platform was used for all sequence-handling tasks in this study.

### Simulation systems

#### SHANK3 SPN-ARR

System S1 is an atomistic model of the WT SPN-ARR domain (residues 2–347) of SHANK3. The model is based on the X-ray structure of the N-terminal domains of SHANK3 (PDB:5G4X) ^14^. System S2 comprises a similar model where the residue N52 of the 5G4X structure is mutated to arginine. To complement this, in System S3 we constructed the N52R mutant from the coordinates of the X-ray structure of SHANK3–Rap1A (PDB:6KYK) (Cai et al., 2019), but without Rap1A proteins. Together these systems served to study the structural dynamics of the SPN-ARR domain in a water environment.

#### SHANK3 SPN-ARR with Rap1A

System S4 entails a SHANK3 SPN-ARR domain (residues 5–363) complexed with two GNP-loaded Rap1A proteins (residues 1–166). The complex was extracted from the SHANK3–Rap1A structure (PDB:6KYK) (Cai et al., 2019). To expedite conformational sampling, SHANK3 was mutated to the N52R form, which in simulations of System 2 was observed to undergo structural opening. The Rap1A-bound SPN-ARR constructs in System S4 were compared to Systems S1-3 to shed light on the role of Rap1A in the dynamics of the SHANK3 N-terminal domains.

#### Free energy of opening in SHANK3 SPN-ARR

In Systems S5 and S6, we elucidated the free energy of SPN-ARR opening in the WT and N52R mutant systems, respectively. To this end, we used a series of umbrella sampling simulations where we sampled the opening of the SPN-ARR structure, using the distance between these two domains as the reaction coordinate. The simulation parameters of the systems (S1-S6) are described below.

### Simulation models

Simulation models were built using the CHARMM-GUI portal ^70,71^. Accordingly, all the mutations and post-translational modifications were implemented with CHARMM-GUI ^70^. Interactions were described by the all-atom CHARMM36m force field ^72^. Water molecules were described by the TIP3P water model ^73^. Potassium and chloride ions described by the CHARMM36m force field were added to neutralize the charge of the systems and to reach the physiological saline concentration (150 mM).

### Simulation parameters

We used the GROMACS simulation software package (version 2018) to run the simulations ^74^. Initiation of the simulation runs followed the general CHARMM-GUI protocol: the simulation systems were first energy-minimized and then equilibrated with position restraints acting on the solute atoms ^72^. Key parameters of production simulations are described in Table 1. We used the leap-frog integrator with a timestep of 2 fs to propagate the simulations ^75^. Periodic boundary conditions were applied in all three dimensions. Atomic neighbors were tracked using the Verlet lists, and bonds were constrained by the LINCS algorithm ^76^. Lennard-Jones interactions were cut off at 1.2 nm, while electrostatic interactions were calculated using the smooth particle-mesh Ewald (PME) algorithm. The pressure of the system was set to 1 bar and coupled isotropically using the Parrinello-Rahman barostat with a time constant of 5 ps ^77^. Temperature was set to 310 K and coupled separately for solute and solvent atoms using the Nosé—Hoover thermostat with a time constraint of 1 ps. Simulation trajectories were saved every 100 ps. Random initial velocities were assigned for the atoms from the Boltzmann distribution at the beginning of each simulation. For the remaining parameters, we refer to the GROMACS 2018.8 defaults ^74^.

In the umbrella sampling simulations (systems S5 and S6), we opened the initially closed SPN-ARR structure by pulling the SPN domain away from the ARR domain using a series of umbrella sampling windows (see Table 1). Starting from the closed structure, we increased the SPN-ARR distance by 1.4 Å at a time between consecutive sampling windows. This ensured sufficient overlap between the consecutive windows. Here, we exploited the *pull_init* option of GROMACS to set a new distance for each of the 300 ns windows. All 300 ns per window were used for the analysis of the potential of mean force using the weighted histogram analysis method (WHAM), which is implemented as the *gmx wham* code in GROMACS ^78^. In the sampling windows, a force constant of 1000 kJ mol^−1^ nm^−2^ was used to constrain the SPN domain at each distance from the ARR domain. Meanwhile, the ARR domain was restrained (1000 kJ mol^−1^ nm^−2^) from the heavy atoms of residues 115-137, 154-170, 188-203, 221-237, 255-270, 288-303, and 321-333. These residues were selected because they span the entire length of the ARR domain but do not reside at its SPN binding interface. Error estimates were calculated by bootstrap analysis implemented within the *gmx wham* code.

### Flow cytometry (FACS) to analyze β1-integrin activity

Cell-surface β1-integrin activity was analyzed in transfected CHO cells with a previously described, FACS-based assay ^79^. CHO cells were detached using Hyclone® HyQTase (Thermo Fisher Scientific Inc, #SV300.30.01), and resuspended in warm, serum-free medium. The cells were incubated for 40 minutes in rotation at RT with Alexa Fluor 647-labelled fibronectin 7-10 fragment in the presence or absence of 5 mM EDTA (the negative control). The cells were washed with cold Tyrodes buffer (10 mM Hepes-NaOH pH 7.5, 137 mM NaCl, 2.68 mM KCl, 0.42 mM NaH_2_PO_4_, 1.7 mM MgCl_2_, 11.9 mM NaHCO_3_, 5 mM glucose, 0.1 % BSA) and were fixed with 2 % PFA in PBS for 10 min at RT. The PFA was washed away with cold tyrodes and cells were incubated with an anti-α5-integrin antibody (clone PB1, Developmental Studies Hybridoma Bank) in Tyrodes for 30 min at RT with rotation followed by Alexa Fluor 555-conjugated secondary antibody in rotation for 30 min at RT. Cells were washed twice with Tyrodes and resuspended in PBS. The fluorescence signal was analyzed using LSRFortessa (BD Biosciences, Franklin Lakes, NJ) and analyzed using Flowing Software 2.5.1. Viable single cells were gated based on forward scatter area (FSC-A) and side scatter area (SSC-A). GFP-positive cells were further gated from the total population, and Alexa 647 intensity was measured for each sample. The results were normalized to total α5β1-integrin staining. The α5β1 integrin activation index was defined as AI = (F–F_0_)/(F_integrin_), where F is the geometric mean fluorescence intensity of fibronectin 7-10 binding and F_0_ is the mean fluorescent intensity of fibronectin 7-10 binding in EDTA-containing negative control. F_integrin_ is the normalized average mean fluorescence intensity of total α5β1 integrin (PB1).

### Zebrafish microinjections, plasmids & *in vitro* transcription

To generate templates for mRNA *in vitro* transcription, pHAGE-shank3 plasmids were digested with EcoRI and NotI and the plasmid backbone was isolated on agarose gel. Insert was annealed by using annealing oligonucleotides (−5’-*AATTCGATCGTAATACGACTCACTATAGGGA*-3’) and (5’-*GGCCTCCCTATAGTGAGTCGTATTACGATCG*-3’). Annealed product was ligated into digested vector using T4 DNA ligase (NEB). The ligated plasmid was transformed into DH5α competent bacteria. Plasmids were isolated from bacteria clones with NucleoSpin Plasmid Easypure kit (Macherey-Nagel) and screened using digestion with PvuI enzyme. Correct plasmids were linearized with PvuI and used in HiScribe™ T7 ARCA mRNA Kit (with tailing) (NEB) and purified with RNA-25 Clean & Concentrator RNA purification kit (Zymo Research).

Wild-type (AB strain) zebrafish were housed under license MMM/465/712-93 (issued by the Ministry of Agriculture and Forestry, Finland) and embryos were obtained via natural mating. Right after spawning, the embryos were collected and injected with 3.5 ng of either control morpholino oligo or with *shank3a (*AGAAAGTCTTGCGCTCTCACCTGGA) and/or *shank3b (*AGAAGCATCTCTCGTCACCTGAGGT) targeting morpholino oligos ^55^ into 1-4 cell stage embryos using Nanoject II microinjector (Drummond Scientific). To study the effects of shank3 mutations, in vitro transcribed mRNAs were co-injected into embryos. After injections, the embryos were placed in E3 medium (5 mM NaCl, 0.17 mM KCl, 0.33 mM CaCl_2_, 0.33 mM MgSO_4_) supplemented with pen/strep and incubated at 28.5°C.

### Zebrafish motility assay

To analyse motility of zebrafish embryos, 15 μl of 2mg/ml pronase solution was added a day after injections to facilitate hatching. At two days post fertilization, the embryos were transferred to 96-well plates (1 embryo/well). The motility analysis was carried out at 28.5°C using Daniovision instrument (Noldus IT) by imaging the plate at 30 fps for 60 min. First, a 30 min baseline was followed by three 10 min cycles of light/dark (5 min each). After this, 20 mM pentylenetetrazole (PTZ, Sigma-Aldrich) was added to stimulate motility of embryos and a similar program was run again. The speed and total distance moved was analysed using Ethovison XT software (Noldus IT). The first 20 min of baseline was removed and remaining 40 min was used in statistical analyses. Movements were filtered using 0.2 mm minimum distance filter, to reduce background noise, and a maximum movement filter of 4 mm. Average swim speed, total distance moved and the fraction of time spent moving were quantified.

### Zebrafish eye pigmentation assay

To analyse the effects on zebrafish eye pigmentation, the microinjected embryos of 30 hpf (hours post fertilization) of age were dechorionated using forceps. After dechorionation, embryos were anesthetized using Tricaine (160 mg/ml) and imaged using Zeiss AxioZOOM stereomicroscope. Image analysis was carried out using ImageJ/FIJI. First, the images were inverted and background was removed (radius 50). Then, the eyes were outlined manually with a segmented line selection tool and intensity was measured.

### Data analysis & statistics

Whenever data were deemed to follow a non-normal distribution (according to Shapiro-Wilk normality test), analyses were conducted using non-parametric methods. The names and/or numbers of individual statistical tests, samples and data points are indicated in figure legends. All statistical analyses were performed with GraphPad Prism 7 or 8 software and a P-value 0.05 or less was considered as statistically significant.

