## Extended data figures and video legends for "Conformational dynamics regulate SHANK3 actin and Rap1 binding"

**Supplementary information**

- **Extended data figures 1-5 and associated legends**
- **Supplementary video 1 and 2 legends**

### Extended data figures and legends

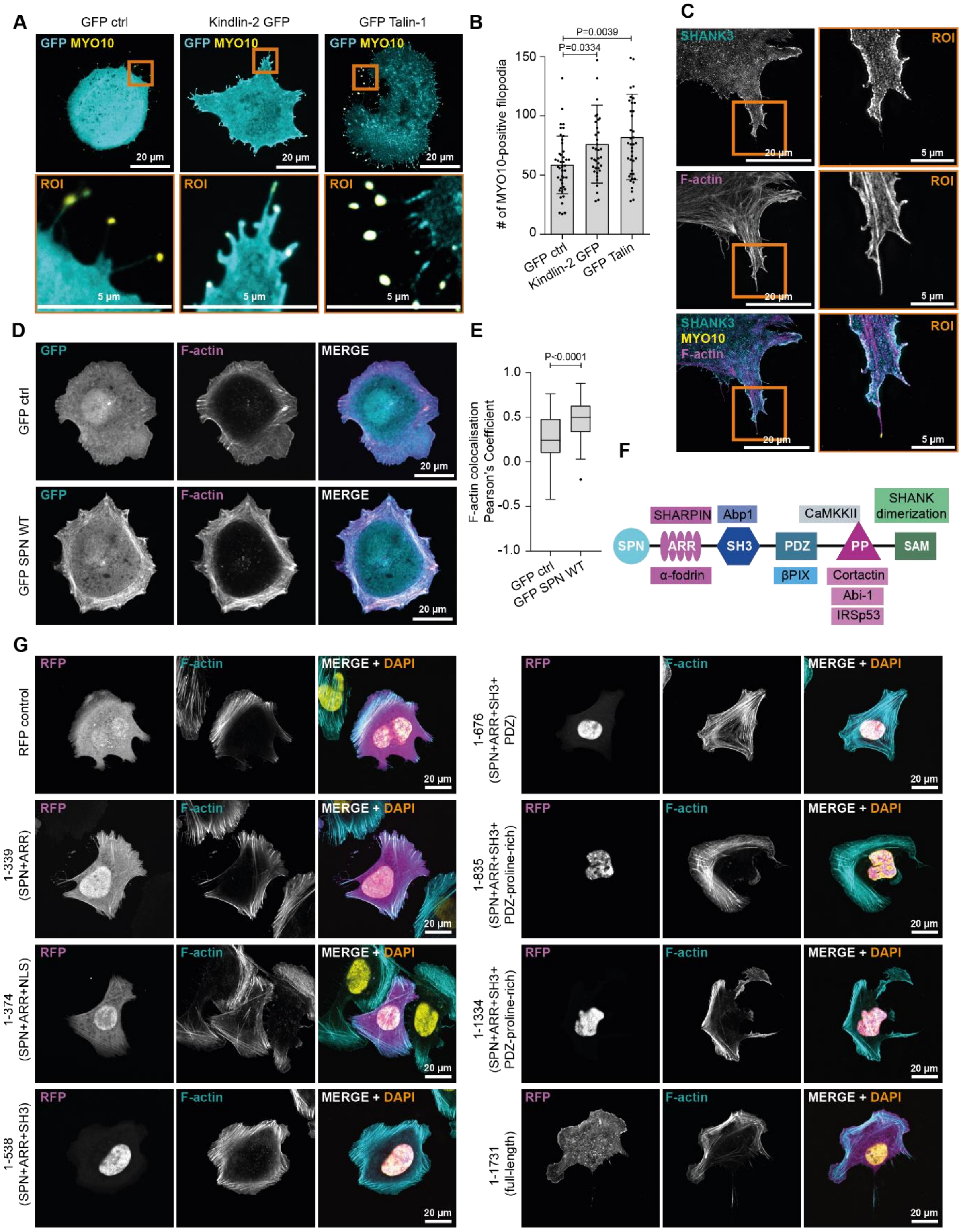


**Extended Data Figure 1. SHANK3 SPN domain colocalization with actin is inhibited in longer SHANK3 fragments. A, B,** Analysis of filopodia formation in U2OS cells co-expressing either GFP control, kindlin-2 GFP or GFP talin together with MYO10-mCherry and plated on fibronectin for 2 h. Representative bottom plane confocal images (A) and quantification of filopodia number (B) are shown. **C,** Analysis of SHANK3 localization along filopodia in U2OS cells co-expressing GFP SHANK3 WT and MYO10-mCherry plated on fibronectin for 2 h and stained for F-actin (SiR-actin). **D, E,** Analysis of F-actin (attophalloidin-647) and GFP colocalization in HEK293 cells expressing either GFP control or GFP SPN and plated on fibronectin for 1 h. Representative bottom plane confocal images (D) and quantification

(E) using the coloc2 ImageJ plugin from one experiment are shown. **F**, Schematic of SHANK3 functional domains and each domain's actin-related binding partners. **G,** Analysis of SHANK3 subcellular localization in U2OS cells expressing different SHANK3-mRFP fragments, plated on fibronectin (3-4 h) and stained for F-actin (attophalloidin-647). Representative bottom plane confocal images from two independent experiments are shown. All representative micrographs and data are from n = three independent experiments unless otherwise indicated. Data are mean ± s.d. (B) or presented as Tukey box plots (E). Statistical analyses: (B) Kruskal-Wallis non-parametric test and Dunn's multiple comparisons post hoc test. (E) Mann-Whitney two-tailed T-test. Number of cells analyzed: (B) 43 cells (GFP ctrl), 38 (Kindlin-2 GFP) and 41 (GFP Talin). (E) 79 (GFP ctrl) and 84 (GFP SPN).

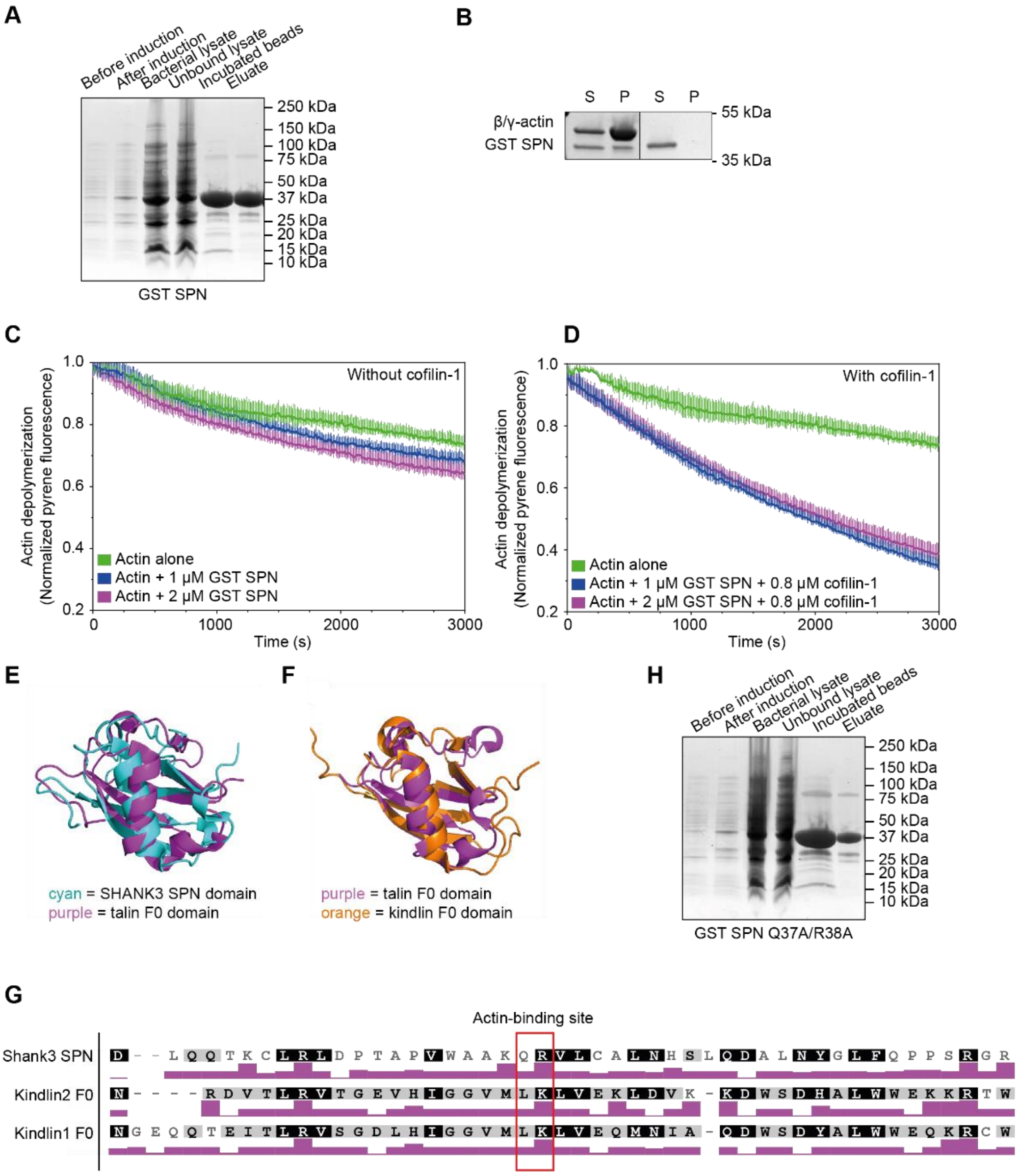


**Extended Data Figure 2. SHANK3 SPN does not affect actin filament stability. A,** Recombinant GST-tagged SPN protein was expressed and purified from *E. coli*. Samples were resolved by SDS-PAGE and visualized by Coomassie Blue staining. Figure shows a representative gel. **B,** Analysis of GST-tagged SPN interaction with β/γ-actin filaments in co-sedimentation assays. S, supernatant fraction; P, pellet fraction; n = 1 experiment. **C, D,** Analysis of spontaneous (C) or cofilin-induced (D) disassembly of β/γ-actin (4 μM of pre-polymerized β/γ-pyrene-actin) filaments in the presence of GST SPN (1 or 2 μM) after a 5 min incubation, monitored by a decrease in pyrene-actin fluorescence. **E, F,** Superimposition of the talin F0 domain (PDB: 2KC1) with either SHANK3 SPN (PDB: 5G4X) (F) or the kindlin F0 domain (PDB: 2KMC). **G,** Sequence alignment between the SHANK3 SPN and the kindlin1/2 F0 domains. The heights of the purple colored bars represent amino acid pI (isoelectric point) values. The values are normalized such that the amino acid with the lowest PI has a value 0 and the highest a value of 1. Other amino acid's values are interpolated to linearly fit this range and shown to highlight similarities in the local charge distribution of the actin binding site residues.

**H,** Recombinant GST-tagged SPN Q37A/R38A protein were expressed and purified from *E. coli*. Samples were resolved by SDS-PAGE and visualized by Coomassie Blue staining. Figure shows a representative gel. All data are from three independent experiments unless otherwise indicated. Error bars represent s.d.

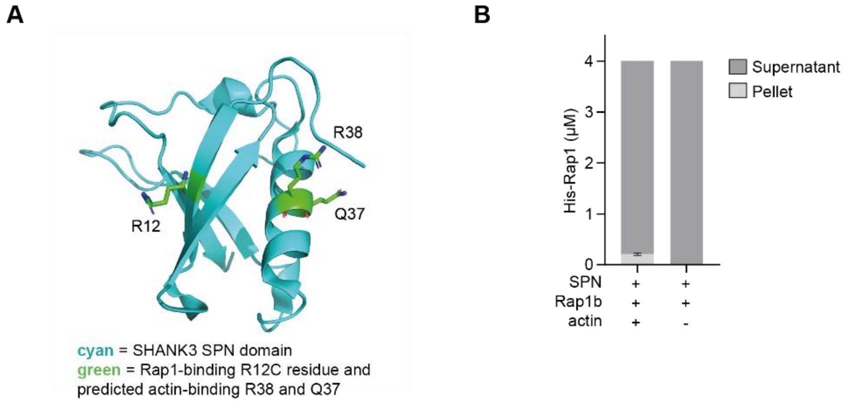


**Extended Data Figure 3. The SHANK3 SPN domain structure and predicted actin binding residues and Rap1 actin co-sedimentation. A,** Visualization of SHANK3 SPN domain (PDB 5G4X) with the Rap1-binding residue R12 and the putative actin-binding residues Q37 and R38 highlighted. **B,** Analysis of active GMPPCP-loaded (GTP-analogue) His-Rap1b (4 μM) interaction with β/γ-actin filaments (12 μM) in co-sedimentation assays after 50 min incubation. The amount of Rap1b in the supernatant and pellet fractions was quantified. His-Rap1b does not interact with actin filaments in the presence of SPN. Five independent experiments; error bars represent s.e.m.

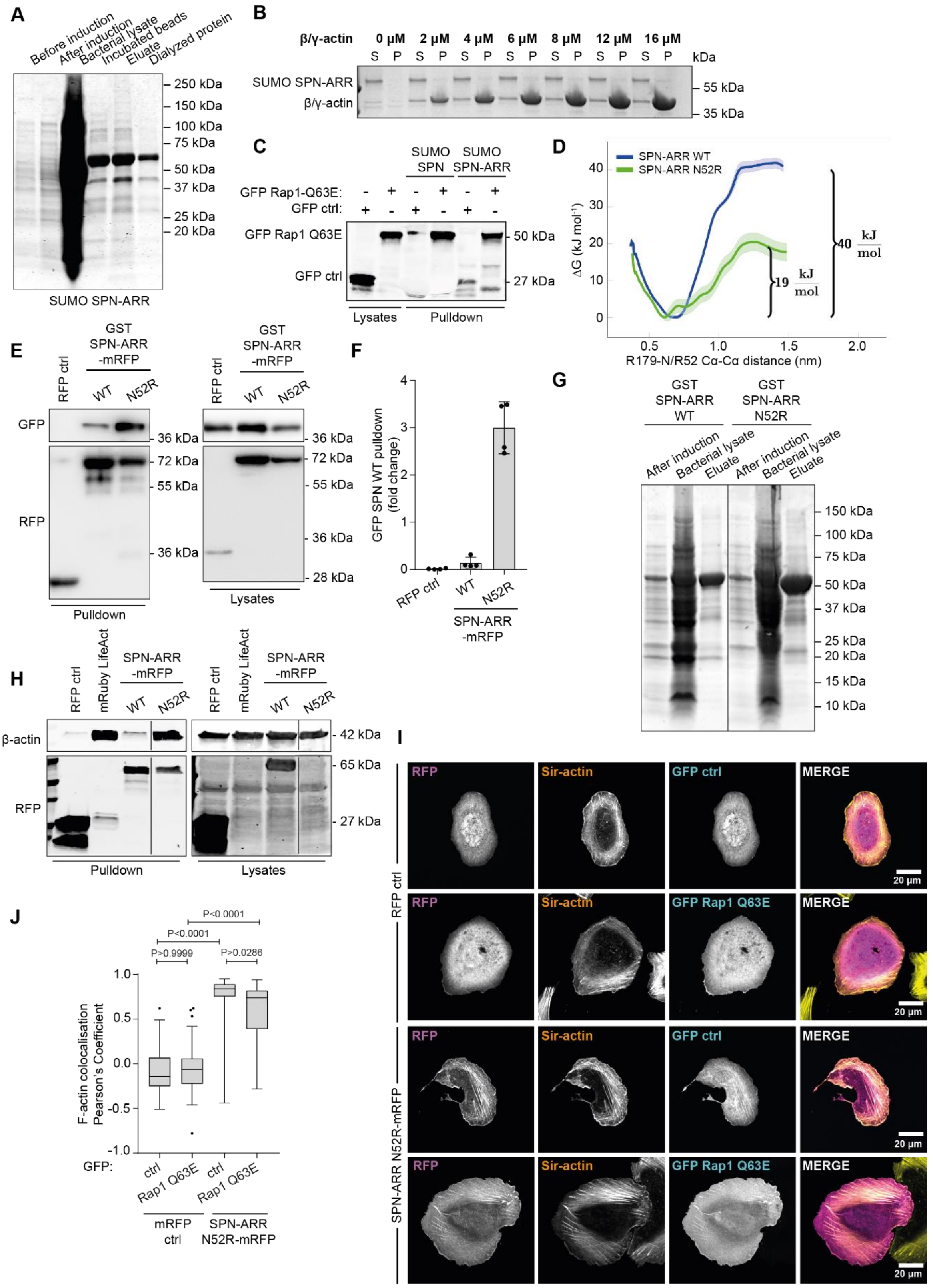


**Extended Data Figure 4. Active Rap1 inhibits SHANK3 actin interaction. A,** Recombinant His$_6$-SUMO SPN-ARR expressed and purified from E. coli. Samples were resolved by SDS-PAGE and visualized by Coomassie Blue staining. Figure shows a representative gel. **B,** Analysis of His$_6$-SUMO SPN-ARR binding to β/γ-actin filaments in co-sedimentation assays. A Coomassie blue stained gel from a representative experiment is shown. **C,** His$_6$-SUMO-tagged SPN or SPN-ARR bound to nickel sepharose beads were used in pulldowns in U2OS cells expressing either GFP (negative control) or constitutively active GFP Rap1 Q63E. Input lysates and IP samples were analyzed using an anti-GFP antibody. **D,** Free energy profiles of the opening of SHANK3 SPN-ARR. The data are calculated through Umbrella Sampling atomistic MD simulations (Systems S5 and S6 in Table 1). The two SHANK3 domains are bound at the distance of 0.6 nm, and in the open conformation at 1.4 nm. **E, F,** Representative RFP-trap pulldown in HEK293 cells co-expressing GFP SPN WT together with either RFP control (negative control), SPN-ARR WT-mRFP or SPN-ARR N52R-mRFP (E) and quantification (F). Input lysates and IP samples were analyzed using RFP and GFP antibodies as indicated. Data are representative of four independent experiments. **G,** Recombinant GST SPN-ARR WT and N52R expressed and purified from E. coli. Samples were resolved by SDS-PAGE and visualized by Coomassie Blue staining. Figure shows a representative gel. **H,** RFP-trap pulldown in HEK293 cells expressing either RFP control (negative control), mRuby-LifeAct (positive control), SPN-ARR WT-mRFP or SPN-ARR N52R-mRFP. Input lysates and IP samples were analyzed using β-actin and RFP antibodies as indicated. **I, J,** Analysis of F-actin (SiR-actin) and RFP colocalization in U2OS cells co-expressing RFP control or SPN-ARR N52R-mRFP together with either GFP control or GFP Rap1 Q63E. Cells were plated on fibronectin-coated glass-bottom dishes (3-4 h). Representative bottom plane confocal images (I) and quantification (J) using the coloc2 ImageJ plugin are shown. All data are from three independent experiments unless otherwise indicated. Data are mean ± s.d. (F) or displayed as Tukey box plots (J). Number of cells: (F) 63 (RFP ctrl+GFP ctrl), 65 (RFP ctrl+GFP Rap1 Q63E), 68 (SPN-ARR N52R+GFP ctrl) and 64 (SPN-ARR N52R+GFP Rap1 Q63E). Statistical analysis: (E) Kruskal-Wallis non-parametric test and Dunn's multiple comparisons post hoc test.

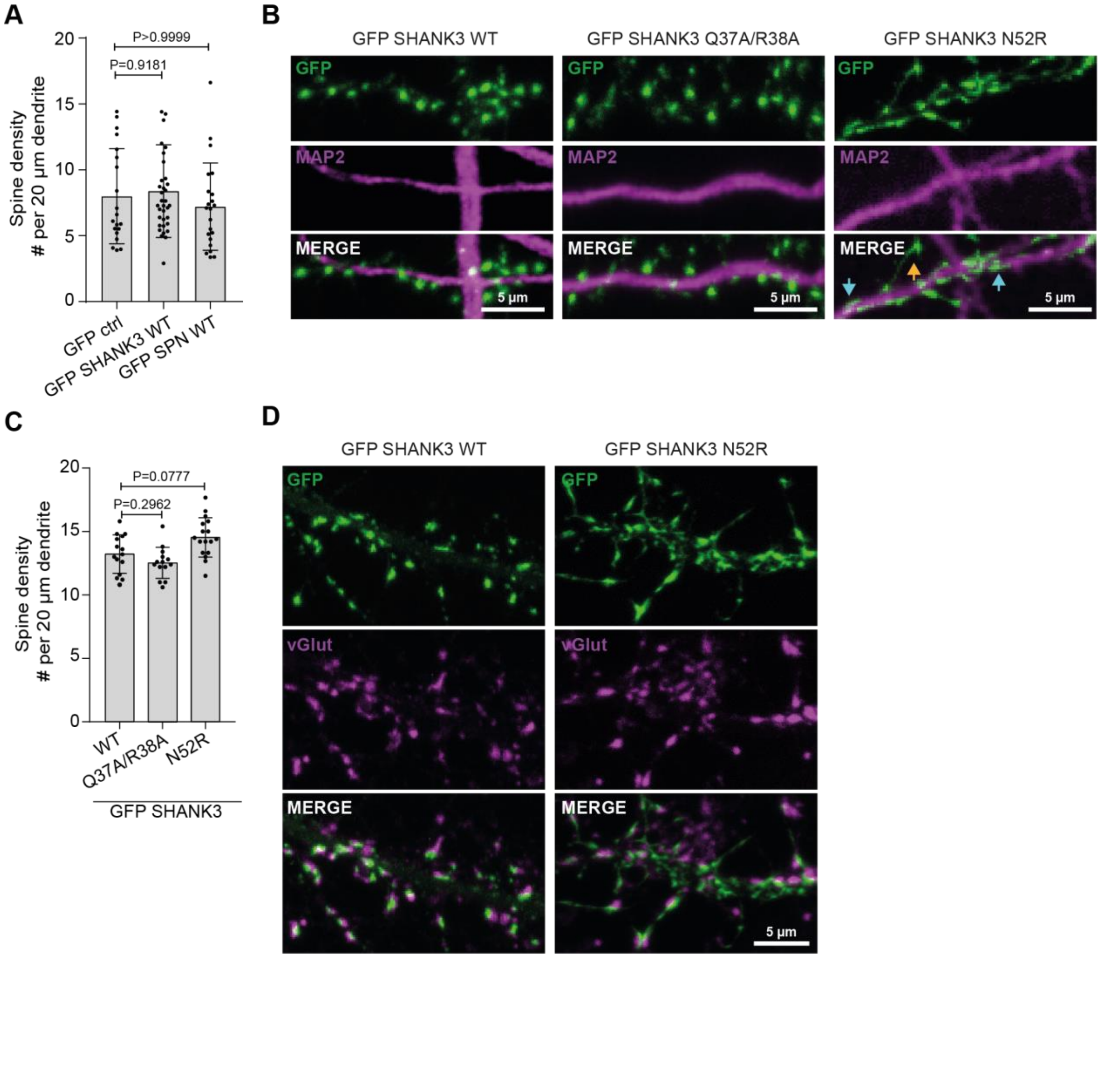

**Extended Data Figure 5. The effects of GFP SHANK3 mutants and GFP SPN in WT primary neurons. A,** Quantifications of spine density of WT primary rat hippocampal neurons expressing the indicated constructs and fixed at DIV16-18. Representative maximum intensity projection confocal images shown in Fig. 6B. **B-D,** Representative maximum intensity projection confocal images (B, D) and quantification (C) of WT primary rat hippocampal neurons expressing the indicated constructs and fixed at DIV16-18. The neurons were stained with the dendritic marker MAP2 (microtubule-associated protein 2) (B) or vesicular glutamate transporter (vGlut) (D). Orange arrow highlights thin spines and blue arrows highlight stubby spines (quantification shown in Fig. 6D). Data represent mean ± s.d : spine density in secondary dendrites; n = 20 (GFP ctrl), 35 (GFP SHANK3 WT) and 22 (GFP SPN WT) neurons (A) and in number of dendrite branches: 45 from 15 neurons (C). Statistical analysis: (A, C) Kruskal-Wallis non-parametric test and Dunn's multiple comparisons post hoc test.

### Supplementary video legends

**Supplementary Video 1.** Atomistic MD simulation of SHANK3 SPN-ARR as both the wild-type and N52R mutant (Systems S1 and S2 in Table 1). The ARR domains are colored orange while the SPN domains are depicted in cyan. Amino acid residues of the ARR domain that are within 0.3 nm from residue 52 are highlighted with licorice representation.

**Supplementary Video 2.** Atomistic MD simulation of the N52R mutant of SHANK3 SPN-ARR bound to two Rap1 proteins (System S4 in Table 1). The ARR domains are colored orange while the SPN domains are depicted in cyan. Rap1 molecules are colored with shades of green. Residues R52 and R179 are highlighted with blue beads.
